# Continuous generation of short-lived resident effector CD8 T cells sustains alopecia areata

**DOI:** 10.64898/2026.11.16.766888

**Authors:** Yoonji Lee, Junghwan Choi, Myeong-seok Lee, Soonjoung Kim, Jin Joo Kang, Suhjeon Shin, Tae Hyun Kim, Sang Hyeok Ji, Suyeong Park, Sihyung Joo, Kunhee Lee, Cheol Lee, Hanjae Lee, Se Jin Im, Ohsang Kwon, June-Yong Lee, You Jeong Lee

## Abstract

CD8 T cells mediating chronic tissue inflammation have been thought to depend on stem-like progenitors that self-renew and replenish cytotoxic effector cells or exhausted progeny. Here, we show that stem-independent, short-lived resident effector CD8 T (T_RE_) cells sustain alopecia areata (AA), a common autoimmune disease that destroys hair follicles. AA-T_RE_ cells exhibit TCR-dependent cytotoxicity, undergo rapid apoptosis in the skin, and lack memory or stem-like properties. Following adoptive transfer, donor T_RE_ cells serve only as an initial trigger, whereas the host-derived CD103⁺CD101⁺ pathogenic pool is rebuilt continuously in skin-draining lymph nodes (sdLNs). Single-cell multiome and CUT&Tag profiling revealed that AA-T_RE_ cells epigenetically silence *Tox* and upregulate RUNX2 to initiate the T_RE_ differentiation program in sdLNs, while RUNX3 and AP-1s sustain it in the skin. Notably, CD8 T cells from patients with AA share this conserved regulatory network. Altogether, these findings define a distinct CD8 T cell differentiation state, offering new therapeutic implications for autoimmune diseases.

**Summary:** In alopecia areata, autoreactive CD8 T cells adopt a resident effector fate, in which TOX-independent TCR-driven activation generates short-lived effectors without stem-like progenitors. RUNX2 initiates this program in skin-draining lymph nodes, and RUNX3 and AP-1s sustain it in the skin.

## INTRODUCTION

In cancer and chronic viral infection, persistent antigen exposure drives a TOX-dependent exhaustion program in which stem-like progenitor-exhausted CD8 T cells (T_EX-PROG_) continuously generate terminally exhausted progeny (T_EX-TERM_) (Belk et al., 2022). Autoreactive CD8 T cells in autoimmunity, however, can remain highly cytotoxic despite chronic stimulation. In type 1 diabetes (T1D), autoimmune progenitors (APs) give rise to autoimmune mediators (AMs), which are short-lived effector cells that minimally express TOX (Gearty et al., 2022). In lupus nephritis, stem-like CD8 T cells generate effector progeny that maintain cytotoxicity through IL-15 and IL-21 signaling, albeit with high expression of TOX and PD-1 (Al Souz et al., 2026). Although these effector states differ, both systems rely on a durable progenitor compartment that continually supplies pathogenic effectors. Whether chronic autoimmune tissue injury can instead be maintained by repeated generation of terminal effectors without persistence of the progenitor pool remains unknown.

Alopecia areata (AA) is a prevalent autoimmune disease characterized by inflammatory hair loss driven by autoreactive CD8 T cells (Gilhar et al., 2012). Approximately 2% of individuals are affected over a lifetime, with 14∼25% of patients progressing to total permanent hair loss (Pratt et al., 2017). Although JAK–STAT inhibitors have recently received FDA approval, patients invariably relapse following treatment discontinuation (Dillon, 2021; Xing et al., 2014). The mechanisms by which pathogenic CD8 T cells in AA are activated remain controversial. One model holds that AA is mediated by virtual memory T (T_VM_) cells acting through antigen-independent, innate-like cytotoxicity, in which T cell receptor (TCR) specificity is largely dispensable (Seok et al., 2023). By contrast, clonally expanded CD8 T cells accumulate around hair follicles in both C3H/HeJ mice and human patients (de Jong et al., 2018), and defined TCRs have been implicated in disease progression (Dai et al., 2025), supporting an antigen-driven process. It therefore remains unresolved whether the pathogenic pool is maintained by antigen-independent cytokine-driven activation or TCR-dependent clonal selection, whether disease-sustaining cells persist or are continuously regenerated, and where their terminal tissue program is specified.

Here we address these questions in the C3H/HeJ mouse model of AA. We identify pathogenic CD103⁺CD101⁺ resident effector T (T_RE_) cells that are tissue-resident, terminally differentiated, and short-lived. Transferred T_RE_ cells disappear within three weeks even in lymphopenic hosts, whereas host-derived T_RE_ cells accumulate first in the skin-draining lymph node (sdLN), indicating that the pathogenic T_RE_ pool is maintained by repeated generation rather than persistence or self-renewal of terminal T_RE_ cells. Paired single-cell transcriptomic and TCR analyses, together with TCR ablation, establish clonal selection and TCR dependence, while no canonical stem-like compartment feeding the T_RE_ pool was detected. Multiome and CUT&Tag profiling identify RUNX2 as an early regulator that engages the T_RE_ program in the sdLN, with RUNX3 recruitment and AP-1 activity becoming prominent after skin entry. Human AA CD8 T cells share key transcriptional and chromatin features of this program. These findings define a terminal autoimmune resident-effector fate whose chronic pathogenicity depends on continuous production in the draining lymph node rather than longevity of the effector pool.

## RESULTS

### CD103⁺CD101⁺ resident effector CD8 T cells define the pathogenic AA compartment

C3H/HeJ mice develop AA spontaneously at about 10% incidence (Pratt et al., 2017), and disease can be serially transferred by intradermal injection of CD8 T cells from the sdLN and skin of affected mice into unaffected (UA) hosts (Wang et al., 2015; Xing et al., 2014). NKG2D⁺ CD8 T cells carry this pathogenicity (Xing et al., 2014), and we previously showed that they express CD103 and distribute to the skin, sdLN, liver, and lungs (Lee et al., 2022). Because neither NKG2D nor CD103 is specific to AA, we sought a phenotype that more selectively identifies the disease-associated population. We found that coexpression of CD103 and CD101 distinguished a CD8 T cell population strongly associated with AA across tissues (Fig. 1A, B). We designated these CD103^+^CD101^+^ CD8 T cells as resident effector T (T_RE_) cells for their tissue residency and limited persistence, as established below. T_RE_ cells in AA (AA-T_RE_ cells) overlap with the previously described CD44 super-high (CD44^s-hi^) CD49d^low^ population (Seok et al., 2023), but that phenotype is also present in the liver of UA mice (Fig. S1); therefore, we defined AA-T_RE_ cells as CD103^+^CD101^+^ CD8 T cells throughout this study. Because CD103 expression alone does not establish tissue residency (Gavil et al., 2023), we assessed residency directly by parabiosis. We generated CD45.1 congenic C3H/HeJ mice by backcrossing and surgically joined CD45.1 and CD45.2 AA partners. Most AA-T_RE_ cells failed to equilibrate between partners (Fig. 1C), and limited exchange was similarly observed in CD45.1 UA and CD45.2 AA pairs (Fig. 1D). Thus, AA-T_RE_ cells are predominantly tissue resident. Functionally, AA-T_RE_ cells, but not non-AA-T_RE_ cells isolated from the same AA donors, transferred disease to UA hosts (Fig. 1E), localizing transferable pathogenic activity to the AA-T_RE_ compartment.

**Figure 1.**
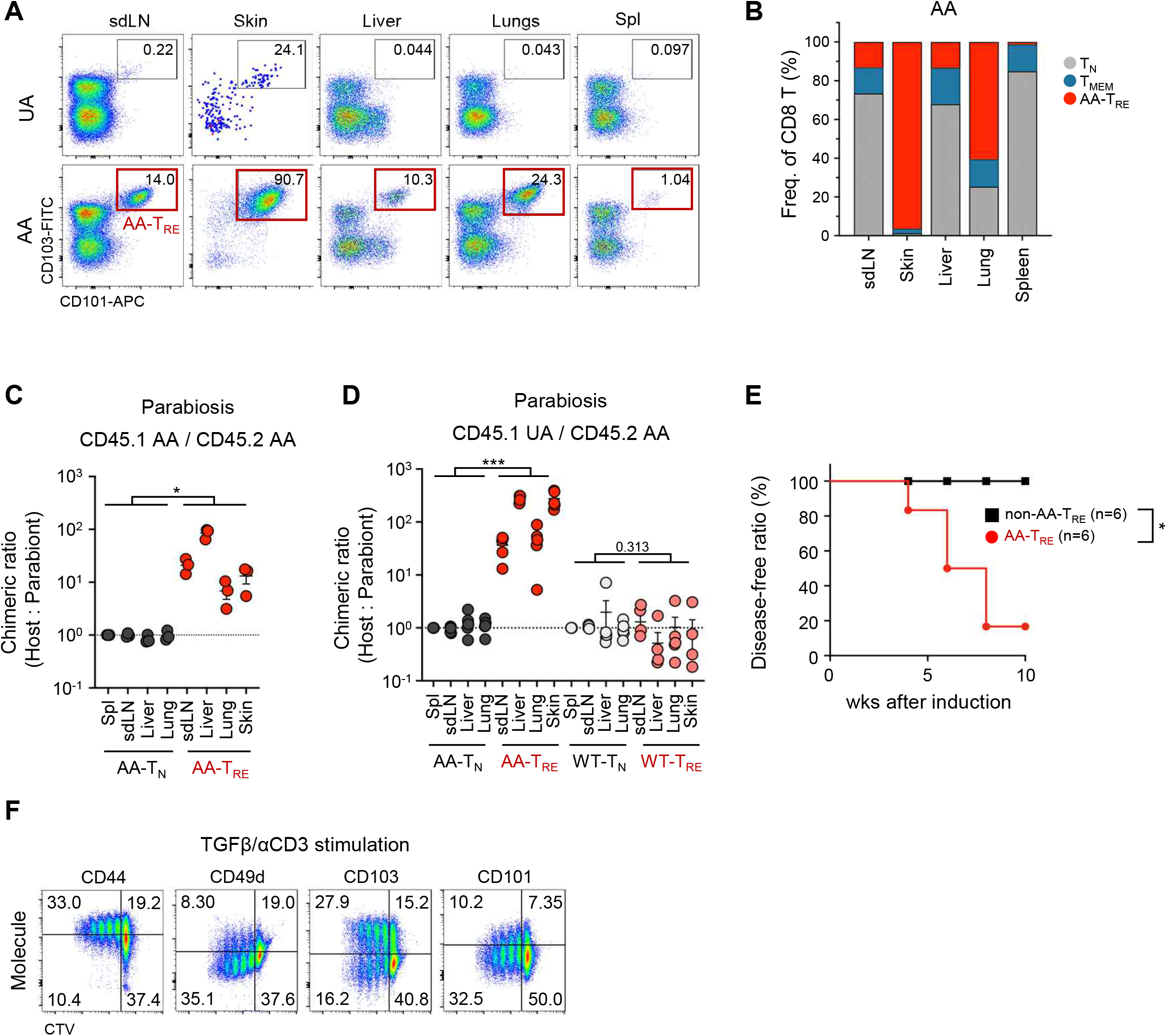
CD103⁺CD101⁺ resident effector CD8 T cells define the pathogenic AA compartment. **(A-B)** Dot plots showing CD103^+^ CD101^+^ CD8 T cells in UA (top row) and AA (bottom row) C3H/HeJ mice (A). Representative data from at least ten independent experiments are shown. Graph showing the proportions of T_N_, T_MEM_, and T_RE_ cells in the indicated organs of AA mice (n=5) (B). **(C-D)** Mice were surgically joined and analyzed two weeks later. Graphs showing the chimeric ratio of host- and partner-derived cells in parabiosis experiments between CD45.1 and CD45.2 AA mice (n=3) (C) and CD45.1 UA and CD45.2 AA mice (n=5) (D). **(E)** Graph showing AA incidence in host mice after intradermal transfer of proportional numbers of non-AA-T_RE_ or AA-T_RE_ cells isolated from AA mice (n=6 each). **(F)** CD44^hi^ CD8 T cells were sorted from the sdLN of UA C3H/HeJ mice, labeled with CTV, and cultured in the presence of TGFβ and anti-CD3/CD28 Dynabeads for 3 days. Representative dot plots from the 3 independent experiments are shown. Each dot represents an individual mouse. Numbers indicate frequencies of cells in each quadrant or adjacent region. Data are mean ± S.D. For statistical analysis, the mean chimeric ratio of the naïve and T_RE_ populations was calculated for each independent experiment, and paired t-tests were performed on these means (C and D). AA incidence curves were analyzed using the Kaplan-Meier method and compared using the log-rank (Mantel-Cox) test (E). *p<0.05, **p<0.01, ***p<0.001. T_N_, naïve T; T_MEM_, memory T; T_RE_, resident effector T; AA, alopecia areata; UA, unaffected.

Because AA-T_RE_ cells phenotypically resemble resident memory T (T_RM_) cells, we asked whether canonical residency cues could recapitulate the T_RE_ phenotype. Anti-CD3/CD28 stimulation in the presence of TGFβ, with or without IL-15, generated CD103⁺CD101⁺CD44^s-hi^CD49d^low^ cells *in vitro* (Fig. 1F and S2), conditions related to those that promote T_RM_ differentiation (Mackay et al., 2013). By contrast, IL-12, IL-15, and IL-18 induced the CD44^s-hi^ phenotype, as previously reported (Seok et al., 2023), but did not induce CD101 or CD103 (Fig. S2). Thus, combined TCR and TGFβ signals are sufficient to reproduce key AA-T_RE_ surface features *in vitro*, linking this pathogenic population to a residency-associated differentiation program without establishing a durable memory fate.

### The AA-T_RE_ pool is maintained by continuous regeneration rather than persistence

Because AA-T_RE_ cells were resident (Fig. 1C, D) yet sufficient to transfer disease (Fig. 1E), we asked whether transferred pathogenic cells persist in recipients or instead initiate a process that recruits a new host-derived population. We transferred CD8 T cells from CD45.1/2 spontaneous AA (SpAA) donors into CD45.2/2 UA hosts and analyzed recipients after disease developed (Fig. 2A). Strikingly, more than 99% of CD8 T cells recovered from the skin, sdLN, liver, lungs, and spleen were host-derived, whereas donor cells were nearly undetectable (Fig. 2B). Kinetic analysis showed that donor AA-T_RE_ cells were lost within 3 weeks, while host-derived AA-T_RE_ cells accumulated first in the sdLN and subsequently in the skin, preceding visible hair loss (Fig. 2C). Consistent with preferential generation or expansion in the sdLN, AA-T_RE_ cells incorporated BrdU most efficiently in the sdLN, particularly during lesion progression (Fig. 2D, E).

**Figure 2.**
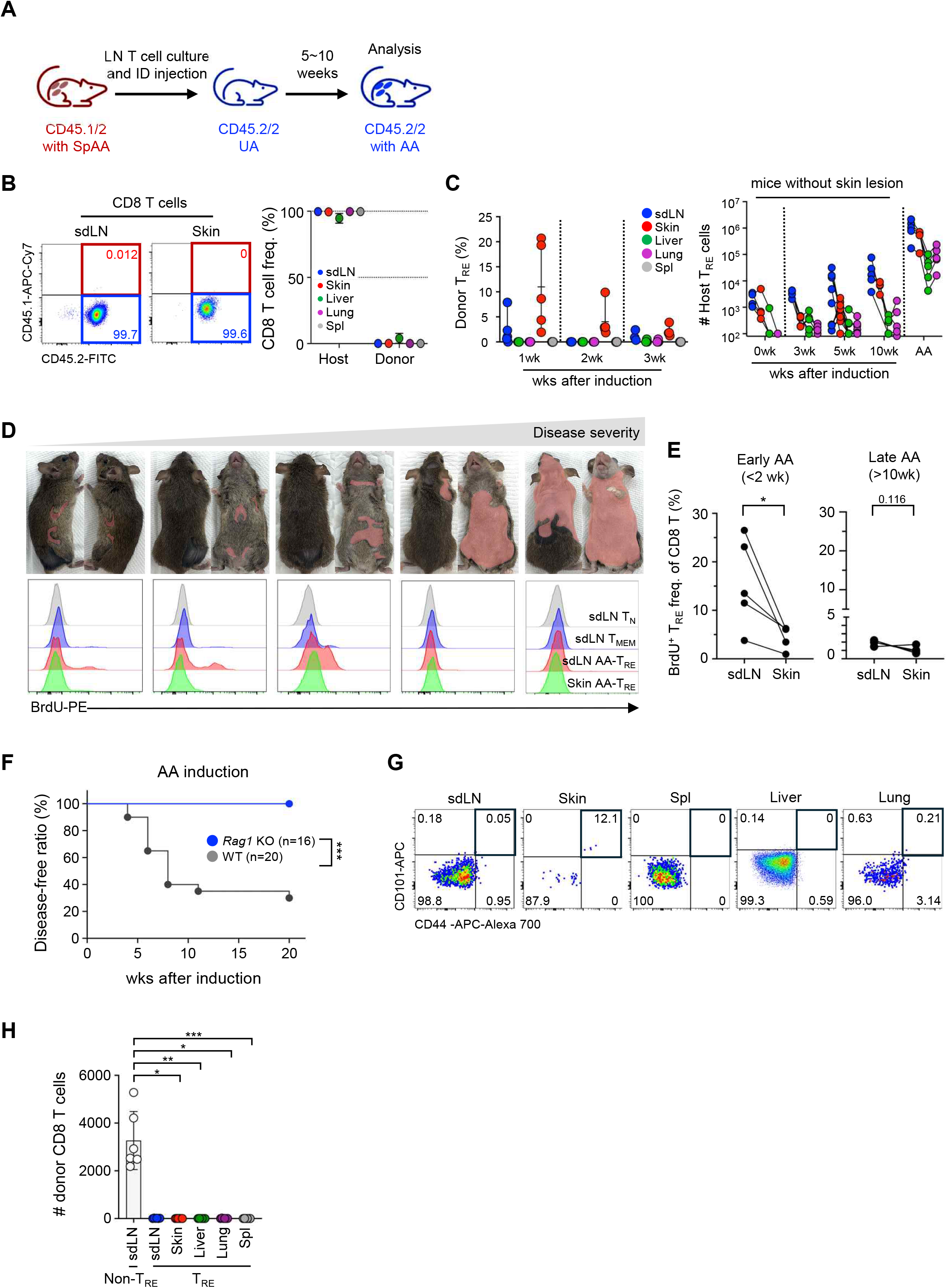
The pathogenic AA-T_RE_ pool is maintained by continuous regeneration rather than persistence. **(A-B)** CD8 T cells from CD45.1/2 SpAA mice were transferred into CD45.2/2 UA hosts intradermally and analyzed after AA development. Schematic figure (A). Representative dot plots (left) and a graph (right) showing the frequencies of donor and host CD8 T cells in the indicated organs (B). Representative data of two independent experiments are shown. **(C)** CD8 T cells isolated from AA mice were transferred into UA hosts as in (A) and analyzed before AA progression. Graphs show the frequencies of donor-derived AA-T_RE_ cells (left) and the number of host-derived AA-T_RE_ cells (right) in the indicated organs at the indicated time points (n=5 at each time point). **(D-E)** *In vivo* BrdU incorporation assay. Images are presented in order of increasing disease severity (top row), and histograms show BrdU incorporation in the indicated populations (bottom row) (D). Graph showing the frequencies of BrdU^+^ AA-T_RE_ cells in sdLN and skin at the indicated time points (E). **(F-H)** WT (n=20) and *Rag1* KO (n=16) C3H/HeJ mice were injected with T cells from AA mice as in (A). Graph showing AA development kinetics (F). Representative dot plots showing donor CD8 T cells in *Rag1* KO C3H/HeJ Host after 10 weeks of T cell transfer (G). Graph showing the frequencies of donor CD8 T cells in Rag KO hosts > 10 wks after adoptive transfer (H). Pooled results from 6 independent experiments are shown. Each dot represents an individual mouse. Numbers indicate frequencies of cells in each quadrant or adjacent region. Data are mean ± S.D. For statistical analysis, a paired t-test (E) and a mixed-effects analysis (REML; repeated-measures ANOVA framework) followed by Dunnett’s multiple comparisons test (H) were performed. AA incidence curves were analyzed using the Kaplan-Meier method and compared using the log-rank (Mantel-Cox) test (F). *p<0.05, **p<0.01, ***p<0.001. T_N_, naïve T; T_MEM_, memory T; T_RE_, resident effector T; SpAA, spontaneous alopecia areata; UA, unaffected.

Donor cell loss could reflect competition with the host repertoire or a cell-intrinsic inability of AA-T_RE_ cells to persist. To distinguish these possibilities, we transferred AA-derived CD8 T cells into *Rag1* KO C3H/HeJ hosts, providing a lymphopenic environment with minimal T cell competition and abundant homeostatic cytokines. Donor AA-T_RE_ cells again failed to persist, and recipients did not develop disease (Fig. 2F, G). Importantly, within the same recipients and from the same inoculum, donor non-T_RE_ CD8 T cells were readily recovered from the sdLN more than 10 weeks after transfer, whereas donor AA-T_RE_ cells were undetectable in every organ examined (Fig. 2H). This selective loss argues that poor persistence is intrinsic to the AA-T_RE_ state rather than a consequence of competition or the transfer procedure. Because *Rag1* KO hosts also lack host CD8 T cells, their resistance to disease is consistent with, but does not independently prove, a requirement for *de novo* host-derived AA-T_RE_ generation.

Together, these data indicate that the pathogenic AA-T_RE_ compartment is not maintained by persistence or self-renewal of the transferred T_RE_ cells. Rather, transferred AA-T_RE_ cells act as short-lived triggers, while a host-derived T_RE_ population emerges first in the sdLN and then populates peripheral tissues. Thus, chronic maintenance of the AA-T_RE_ pool depends on repeated production rather than longevity of the terminal pathogenic effectors, an organization that differs from progenitor-effector systems described in type 1 diabetes and lupus nephritis (Al Souz et al., 2026; Gearty et al., 2022).

### AA-T_RE_ cells terminally differentiate and undergo apoptosis in the skin

We next profiled CD8 T cells across organs of AA mice by 26-plex flow cytometry (Fig. S3A), with additional assessment of T-bet^Zsgreen^, CD122, TGFβRII, TIGIT, and TIM-3 (Fig. 3A-C). t-SNE analysis resolved 16 distinct clusters (c0–15) comprising naïve T (T_N_; c0-c4), CD44^hi^ memory T (T_MEM_; c8-c11), and AA-T_RE_ cells in the skin (c14-c15), sdLN (c13), and liver and lungs (c12), with AA-T_RE_ clusters grouping together by phenotypic similarity (Fig. 3A and S3B-C). Across organs, AA-T_RE_ cells expressed cytotoxicity- and residency-associated molecules, including NKG2D, CD103, CD69, CD49a, and TGFβRII, and were uniformly T-bet-high and Eomes-low (Fig. 3B-C). At the same time, they downregulated CD122, distinguishing the AA-T_RE_ phenotype from conventional memory populations and suggesting progressive acquisition of an activated resident-effector state.

**Figure 3.**
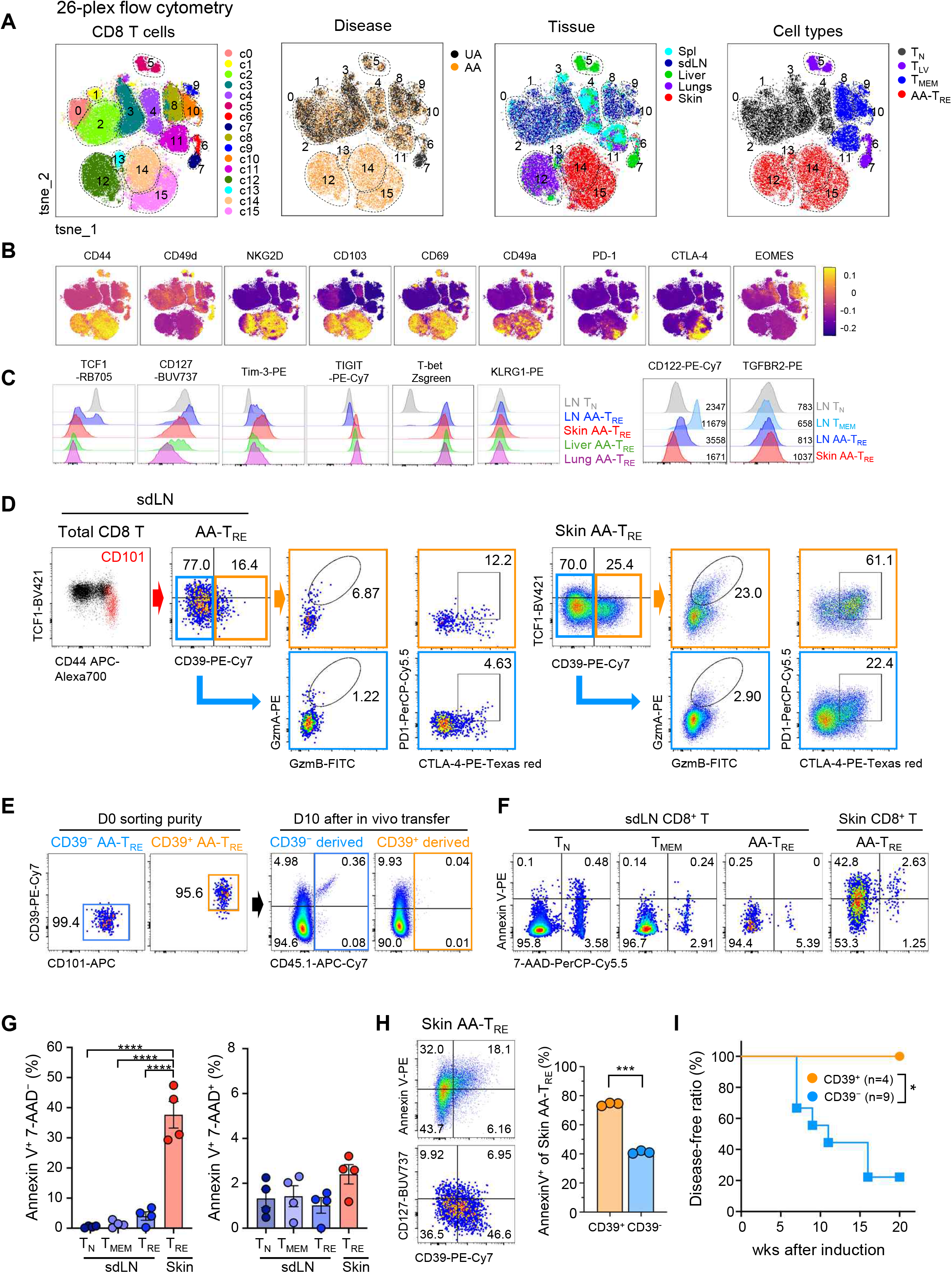
AA-T_RE_ cells terminally differentiate and undergo apoptosis in the skin. **(A-C)** Total CD8 T cells isolated from UA and AA C3H/HeJ mice were analyzed by flow cytometry. UMAP plots showing the indicated disease state, organs, and cell subsets (A). Expression of indicated markers in CD8 T cells in UMAP plots (B) and representative histograms (C). Representative results of 3 to 5 independent experiments are shown. **(D-E)** Dot plots showing the expression of indicated markers in AA-T_RE_ cells in sdLN and skin (D). CD39⁻ or CD39^+^ AA-T_RE_ cells in the skin from CD45.1^+^ AA mice were adoptively transferred into CD45.2^+^ AA hosts. Representative FACS plots showing sorting purity (left two panels) and recovered cells after 10 days of transfer (right two panels) (E). Representative data from 3 independent experiments are presented. **(F-H)** CD8 T cells from AA mice were stained with Annexin V and 7-AAD. Representative flow cytometry plots of indicated populations (F) and graphs showing statistical analysis (G) (n=4). Indicated marker expression in skin AA-T_RE_ (left) and statistical analysis (right) are presented (H) (n=3). Representative data of three independent experiments are shown. **(I)** Graph showing the AA incidence following the intradermal transfer of equal numbers of CD39^+^ (n=4) or CD39⁻ (n=9) donor skin AA-T_RE_ cells. Representative results of two independent experiments are shown. Each dot represents an individual mouse. Numbers indicate frequencies of cells in each quadrant or adjacent region. Data are mean ± S.D. Statistical analysis was performed using one-way repeated-measures ANOVA followed by Tukey’s multiple-comparisons test (G) and a paired t-test (H). Survival curves were analyzed using the Kaplan-Meier method and compared using the log-rank (Mantel-Cox) test (I). *p<0.05, ****p<0.0001. AA, alopecia areata; UA, unaffected; Spl, spleen; T_LV_, liver T; T_N_, naïve T; T_MEM_, memory T; T_RE_, resident effector T.

AA-T_RE_ cells showed little to no TOX expression (Fig. S3D), as previously reported (Dai et al., 2025), despite expressing TIM-3, PD-1, CTLA-4, and TIGIT (Fig. 3B-C). Importantly, these cells retained robust effector activity, producing IFNγ, TNFα, and IL-2 (Fig. S4A-B) and exhibiting high mitochondrial activity, as indicated by oxygen consumption rate and MitoTracker staining (Fig. S4C-D). They were also KLRG1-negative (Fig. 3C). Thus, checkpoint-receptor expression in AA-T_RE_ cells accompanies preserved effector function and does not correspond to the canonical TOX-dependent exhaustion program described in chronic infection and cancer (Khan et al., 2019; Park et al., 2026).

AA-T_RE_ cells downregulated TCF1 in the sdLN and upregulated CD39 in the skin, together with GZMA, GZMB, CTLA-4, and PD-1 (Fig. 3D). To test a precursor-progeny relationship between CD39⁻ and CD39⁺ AA-T_RE_ cells, we sorted each population and transferred equal numbers of them intradermally into congenic AA hosts. After 10 days, CD39⁻ donor cells predominantly acquired CD39, whereas recovery from the CD39⁺ input was approximately ninefold lower than from the CD39⁻ input (0.04% versus 0.36% of skin CD8 T cells) (Fig. 3E and S4E). Skin AA-T_RE_ cells also showed increased Annexin V, and the CD39⁺ subset downregulated CD127 and failed to transfer AA (Fig. 3F-I). GZMA⁺ and TCF1⁺ cells were spatially segregated within the skin, with GZMA^+^ cells preferentially localizing to the epidermal and intrafollicular regions, and TCF1^+^ cells to the interfollicular regions (Fig. S4F-G). Together with the limited persistence observed after transfer (Fig. 2), these data support a progression from CD39⁻ AA-T_RE_ cells to CD39⁺ terminal effectors that rapidly contract in the skin.

Overall, phenotypic progression, transfer, and apoptosis analyses show that AA-T_RE_ differentiation culminates in a short-lived terminal effector state in the skin.

### AA-T_RE_ differentiation is clonally selected and strictly TCR-dependent

Preferential clonal expansion along the AA-T_RE_ trajectory would support antigen-driven selection and provide a direct test of the cytokine-dominant model proposed for AA (Seok et al., 2023). Although skin CD8 T cells in AA form an effectorness gradient (Lee et al., 2023), clonal relationships among AA-T_RE_ cells across organs have not been defined. We therefore performed paired scRNA-seq and scTCR-seq on CD8 T cells from the skin, lungs, liver, spleen, sdLN, and thymus of two spontaneous AA (SpAA) mice (Fig. S5A). UMAP resolved 11 clusters, with AA-T_RE_ cells from multiple organs concentrated in clusters 7, 8, and 10 (Fig. 4A and S5B). These cells were markedly clonally expanded, and identical TCR clonotypes were present across organs (Fig. 4B-D).

**Figure 4.**
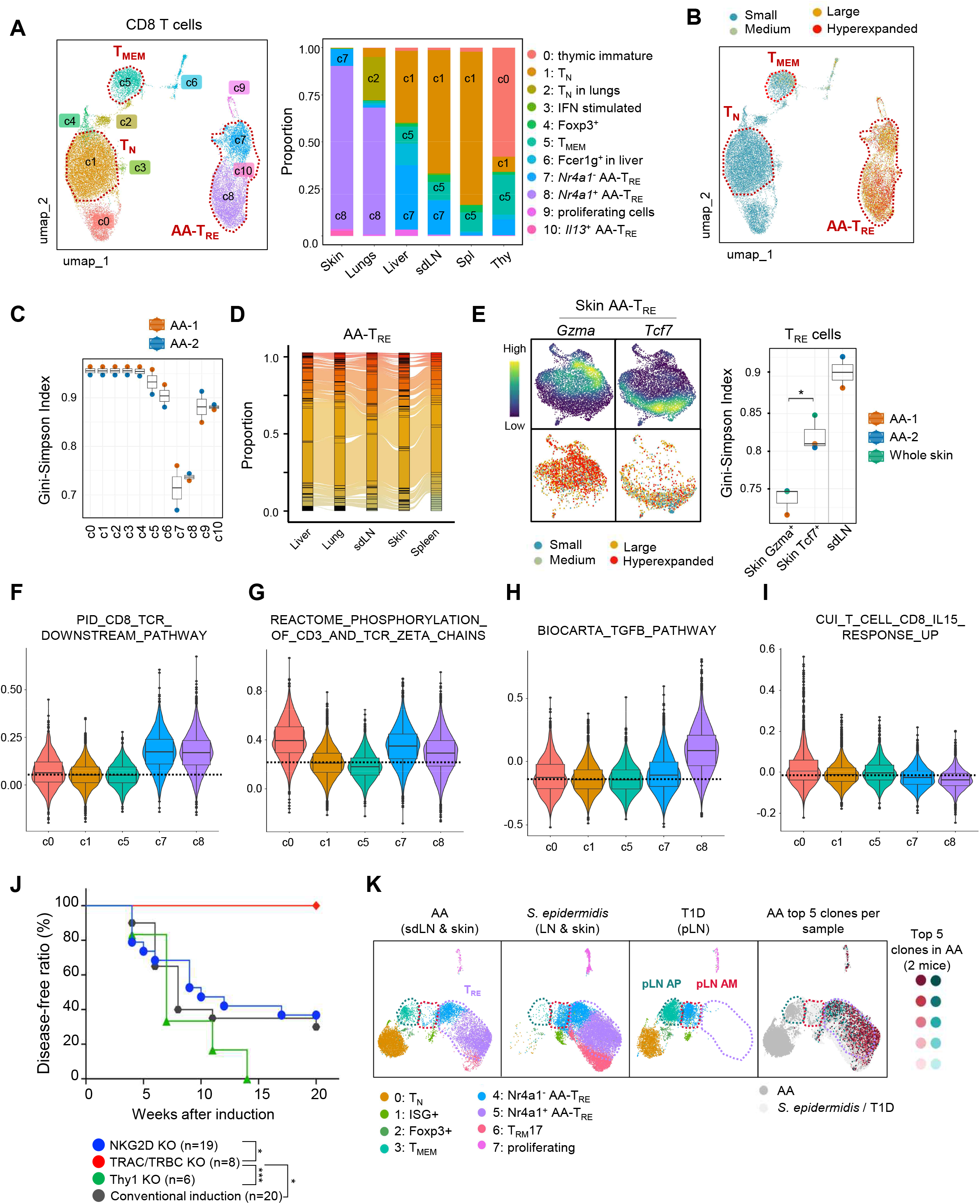
AA-T_RE_ differentiation is clonally selected and strictly TCR-dependent. **(A-I)** CD8 T cells were isolated from indicated organs, and scRNA-seq combined with scTCR-seq was performed. UMAP visualization of scRNA-seq of C3H/HeJ AA CD8 T cells grouped by clusters (left) and fraction of each cell type comprising each organ (right) (A). UMAP displays clonal expansion; small ≤ 0.001, 0.001< medium ≤ 0.01, 0.01 < large ≤ 0.1, 0.1 < hyperexpanded ≤ 1 (B). Gini-Simpson diversity index of CD8 T cells in the indicated clusters (C). Graph showing the overlap of inter-organ clonality in AA-T_RE_ cells (D). Density plots showing *Gzma* and *Tcf7* expression in skin AA-T_RE_ (left, top). UMAPs show *Gzma*^+^ and *Tcf7*^+^ T cells colored by clonal expansion (left, bottom). Graph showing the Gini-Simpson index of the indicated AA-T_RE_ cells (right) (E). Violin plots showing module scores for the indicated gene sets across the specified clusters (F-I). **(J)** Graph showing the disease-free ratio in mice induced with AA by transferring AA sdLN CD8 T cells deleted for the indicated genes using CRISPR-Cas9 (n=6 ∼ 20). Pooled results from 3 independent experiments. **(K)** Integrated UMAP of CD8 T cells from AA skin and sdLNs, public *S. epidermidis*–associated skin and LN datasets (GSE233389), and autoimmune T1D pLNs (GSE151652). UMAPs are split by sample origin (left). The top five expanded clonotypes in each AA sample are highlighted (right). Each dot represents an individual mouse. Data are mean ± S.D. Statistical analysis was performed using an unpaired two-tailed Welch’s t-test (E); Kaplan-Meier curves were compared using the log-rank (Mantel-Cox) test (J); *p<0.05, **p<0.01, ***p<0.001. AM, autoimmune mediator; AP, autoimmune progenitor.

In the skin (c8), *Nr4a1* (Nur77), a direct reporter of TCR signaling that is also expressed in T_RM_ cells (Mackay et al., 2013; Moran et al., 2011), was upregulated (Fig. 4A), and oligoclonal expansion was most prominent among *Gzma*⁺ AA-T_RE_ cells (Fig. 4E, left). Clonal diversity decreased from sdLN AA-T_RE_ cells to skin *Gzma*⁺ AA-T_RE_ cells, consistent with progressive selection of specific clonotypes during differentiation (Fig. 4E, right). TCR-signaling gene sets were also enriched in sdLN and skin AA-T_RE_ cells (Fig. 4F-G and S5C), and this result was reproduced in an independent public AA single-cell dataset (Dai et al., 2025) (Fig. S5D). In parallel, AA-T_RE_ cells showed increased TGFβ signaling and reduced IL-15-associated signaling (Fig. 4H-I), consistent with the TGFβ-dependent phenotype induced *in vitro* and the loss of CD122 observed by flow cytometry. Although AA-T_RE_ cells across organs shared a core transcriptional state, skin AA-T_RE_ cells were transcriptionally distinct from those in liver and lungs (Fig. S5E). These observations support progressive, TCR-associated clonal selection along the sdLN-to-skin AA-T_RE_ trajectory.

We next examined whether TCR expression is required for AA transfer. We deleted *Thy1* (targeting control), *NKG2D* (innate cytotoxicity), or *TRAC/TRBC* in T cells isolated from the sdLN of AA mice using CRISPR–Cas9 and transferred the edited cells intradermally into UA hosts. Deletion of NKG2D or Thy1 did not alter AA incidence, indicating that it is dispensable for disease triggering, although NKG2D marks the pathogenic population (Xing et al., 2014). In contrast, TCR ablation completely prevented AA induction by the transferred cells (Fig. 4J), establishing a requirement for TCR expression in pathogenic activity.

A previous study showed that T cells engineered with clonally expanded AA-associated TCRs can induce disease (Dai et al., 2025). To test whether defined TCRs are sufficient to direct AA-T_RE_ differentiation, we transduced splenic CD8 T cells with expanded AA-associated TCRs and transferred them into UA hosts (Fig. S6A). Upon AA induction, GFP⁺ monoclonal donor cells acquired the CD44^s-hi^CD101⁺ AA-T_RE_ phenotype in both the sdLN and skin (Fig. S6B). In mice that remained disease-free, only small numbers of GFP⁺ donor cells were detected, predominantly in the skin, suggesting tissue tropism without full pathogenic differentiation (Fig. S6C). These data support a role for defined AA-associated TCR specificities in directing T_RE_ differentiation within the appropriate disease context.

Because CD49d^low^ and TOX⁻ phenotypes had been used to support a T_VM_-mediated innate cytotoxicity model of AA (Seok et al., 2023), we asked whether either feature is diagnostic of antigen-independent activation. In two skin-tropic models with defined TCR-driven pathogenicity, melanoma-associated vitiligo (MAV) (Byrne et al., 2011; Frisoli et al., 2020) and 1MOG244-mediated cicatricial alopecia (Alli et al., 2012), pathogenic CD8 T cells also acquired a CD44^s-hi^CD49d^low^ phenotype (Fig. S11). Moreover, pmel-1 CD8 T cells in MAV also showed little TOX expression (Fig. S11C). Thus, CD49d downregulation and low or absent TOX expression are compatible with TCR-driven skin residency and, by themselves, do not establish a T_VM_ origin or antigen-independent pathogenicity.

To contextualize AA-T_RE_ cells across disease settings, we integrated public datasets of skin CD8 T_RM_ cells generated after *S. epidermidis* association (Park et al., 2023) and autoimmune CD8 T cells from T1D (Gearty et al., 2022) (Fig. 4K and S7). AA-T_RE_ cells mapped near *S. epidermidis*-associated skin T_RM_ cells and shared increased *Cd44*, *Cd69*, *Itgae*, and *Tbx21* with reduced *Cd49d* (Fig. S7C). *Il2rb* transcripts remained detectable despite loss of CD122 protein (Fig. 3C), suggesting post-transcriptional regulation of IL-15 responsiveness.

By contrast, T1D CD8 T cells expressed higher *Tox* and *Cd49d* levels and occupied distinct transcriptional regions within the UMAP (Fig. S7C and Fig. 4K). Expanded AA-T_RE_ clonotypes did not map to the regions occupied by the stem-like AP and effector-like AM populations in the T1D dataset (Fig. 4K, far right). These comparisons distinguish the observed AA-T_RE_ state from the canonical T1D progenitor-effector architecture.

Together, TCR editing, defined-TCR transfer, and clonotype tracking establish that AA-T_RE_ differentiation is TCR-dependent and accompanied by progressive clonal selection from the sdLN to the skin. The resulting cells share elements of a skin-residency program but are transcriptionally distinct from the T1D AP/AM architecture and are inconsistent with a purely antigen-independent model of AA pathogenesis (Seok et al., 2023).

### RUNX and AP-1 transcription factors (TFs) are required for AA-T_RE_ differentiation

To identify regulators of AA-T_RE_ differentiation, we performed paired snRNA-seq and scATAC-seq on CD8 T cells isolated from the sdLN and skin of AA mice. Multiome analysis resolved 13 clusters, with clusters 7-10 representing skin AA-T_RE_ populations (Fig. 5A). Compared with T_N_ cells (c0), skin AA-T_RE_ cells (c9) showed increased accessibility at motifs for RUNX2, RUNX3, NUR77, RORA, JUND, and FOSL2 (Fig. S8A-B). SCENIC^+^ analysis likewise identified increased activity of AP-1-family regulons (*Fos, Fosb, Jund,* and *Fosl2*), and RUNX family (*Runx1* and *Runx2*) regulons in AA-T_RE_ cells (Fig. 5B). sdLN AA-T_RE_ cells occupied an intermediate state between sdLN T_MEM_ and skin AA-T_RE_ cells. The inferred gene regulatory network predicted RUNX2 regulation of *Cd101*, *Klrk1*, *Itga1*, and *Nfatc2*, and cFOS regulation of *Cd69*, *Nr4a1*, and *Egr2* (Fig. S8C). Together, these analyses nominated RUNX- and AP-1-family TFs as candidate regulators linking TCR activation, tissue residency, and effector differentiation.

**Figure 5.**
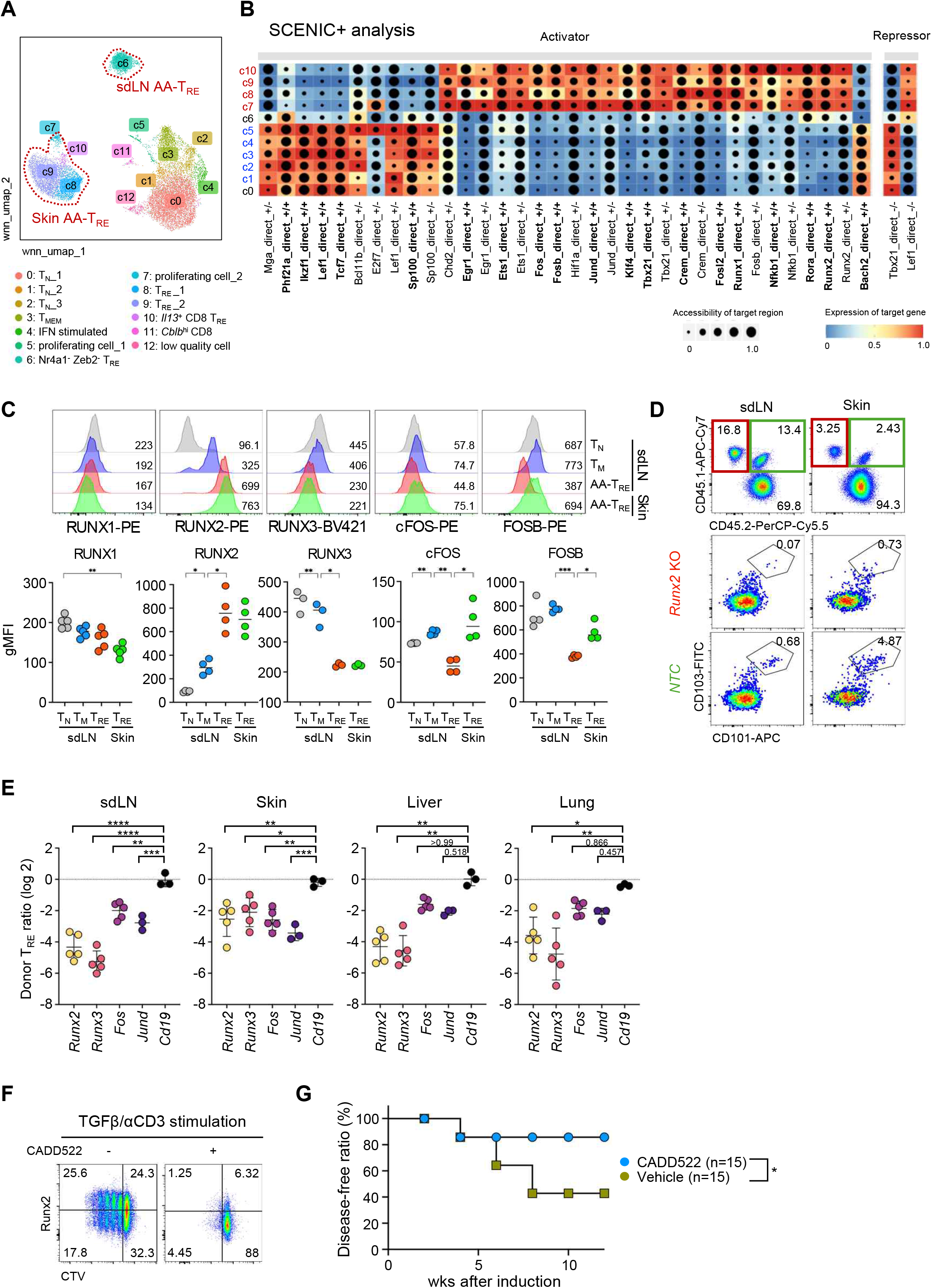
RUNX2, RUNX3, and AP-1 TFs are required for AA-T_RE_ differentiation. **(A-B)** CD8 T cells from the sdLN and skin of C3H/HeJ AA mice were sorted and analyzed by paired snRNA-seq and scATAC-seq. Cells were annotated as in the UMAP in Fig. 3A (A). A dot plot heatmap shows the direct eRegulon derived from SCENIC+ analysis. A color scale shows the expression of the target region, and a size scale shows its accessibility (B). **(C)** Histograms show marker expression in the indicated T cells (top). Graphs show the statistical analysis (bottom) (n=4-5). Pooled results from four independent experiments. **(D-E)** CD8 T cells isolated from UA congenic donors were deleted for RUNX2 (CD45.1/1) or CD19 (CD45.1/2) and transferred into CD45.2/2 AA hosts as depicted in Figure S8D. Representative dot plots show total (top) and donor CD8 T cells in the indicated organs (middle and bottom) (D). Graphs show statistical analysis of the gene-deleted donor AA-T_RE_ ratio in the indicated organs compared with *Cd19* deletion (E). **(F)** CD8 T_MEM_ cells were sorted from the sdLN of UA C3H/HeJ mice, labeled with CTV, and cultured in the presence of indicated cytokines with or without Runx2 blocker (CADD522). Representative data of 3 independent experiments are shown. **(G)** Disease-free ratio in AA-induced mice treated with or without CADD522 once weekly. Each dot represents an individual mouse. Numbers indicate frequencies of cells in each quadrant or adjacent region. Data are mean ± S.D. Statistical analysis was performed using repeated-measures one-way ANOVA followed by Tukey’s multiple comparisons test (C); Kruskal-Wallis test followed by Dunn’s multiple-comparisons test, and the mean value of *Cd19* KO was used as the control for multiple comparisons (E). Kaplan-Meier curves were compared using the log-rank (Mantel-Cox) test (G); *p<0.05, **p<0.005, ***p<0.001.

Although *Runx2* regulon activity was lower in sdLN than in skin AA-T_RE_ cells (Fig. 5B), RUNX2 protein increased progressively from T_N_ through T_MEM_ cells and was already near maximal in sdLN AA-T_RE_ cells (Fig. 5C). In contrast, cFOS and FOSB expression was lowest in sdLN AA-T_RE_ cells and increased after skin entry, whereas RUNX3 protein did not increase along the same transition. These expression and regulon patterns therefore nominated RUNX2 as an early sdLN-associated factor and AP-1 as a later component of the T_RE_ program.

We next tested the functional requirement for these factors. CD8 T cells from UA donors were edited by CRISPR-Cas9 and transferred into congenic AA hosts as depicted in Fig. S8D. Whereas control donor cells generated CD101⁺CD103⁺ AA-T_RE_ cells in the sdLN and skin, *Runx2*-deficient donor cells differentiated into AA-T_RE_ cells 4- to 32-fold less efficiently (Fig. 5D-E). Deletion of *Runx3*, *Fos*, or *Jund* similarly impaired AA-T_RE_ generation (Fig. 5E). In the TCR/TGFβ culture system, RUNX2 was induced, and pharmacologic RUNX2 inhibition with CADD522 markedly reduced T_RE_-like differentiation (Fig. 5F). Consistently, CADD522 treatment also reduced AA induction rates *in vivo* (Fig. 5G).

These functional perturbations establish that RUNX2, RUNX3, Fos, and Jund contribute to AA-T_RE_ generation and prioritize RUNX2 as an early sdLN-associated regulator.

### The resident effector program is licensed by RUNX2 in the sdLN before tissue entry

Because chromatin accessibility alone does not distinguish primed from active regulatory elements, we complemented the multiome analysis with histone CUT&Tag profiling across the AA-T_RE_ trajectory (Fig. 6A-C). H3K4me1 differential enrichment regions (DERs) enriched in skin AA-T_RE_ cells showed a progressive increase from sdLN T_MEM_ through sdLN AA-T_RE_ to skin AA-T_RE_ populations (Fig. 6A), whereas sdLN T_MEM_-enriched regions showed reciprocal dynamics (Fig. S9A-B). Regulatory regions associated with T_RE_-linked genes, including *Runx2*, *Pdcd1*, *Havcr2*, *Cd44*, *Itgae*, and *Cd101*, acquired active or primed enhancer features along this trajectory. Motif analysis of skin AA-T_RE_-enriched H3K4me1 and H3K27ac regions identified RUNX-family and AP-1 motifs (Fig. 6C and S9C-D), supporting stepwise remodeling of the T_RE_ regulatory landscape.

**Figure 6.**
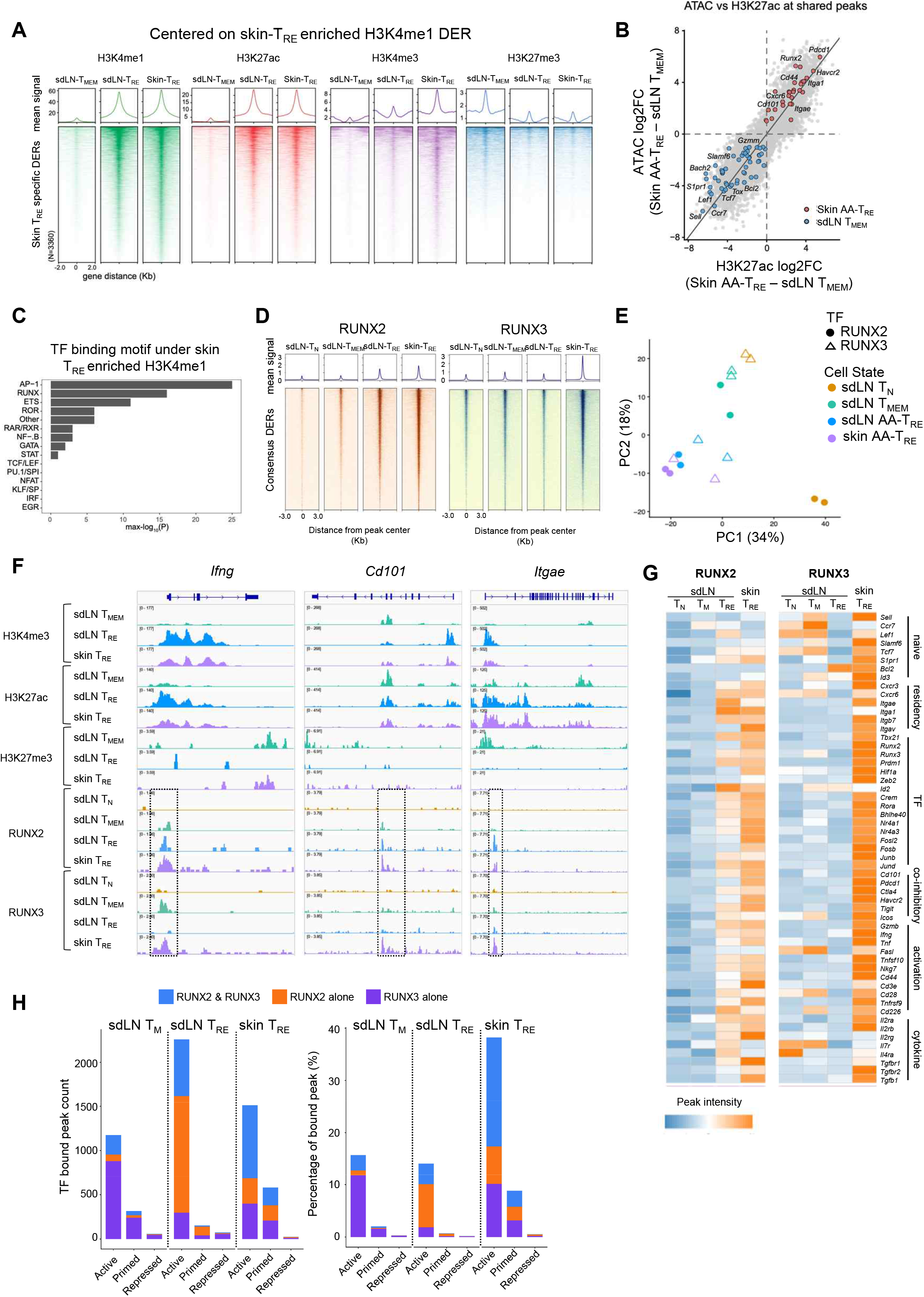
The resident effector program is licensed by RUNX2 in the sdLN before tissue entry. **(A)** Aggregated signal profiles (top) and heatmaps (bottom) show normalized CUT&Tag signals for multiple histone modifications centered on H3K4me1 DERs enriched in skin AA-T_RE_ cells compared to sdLN T_MEM_ cells. **(B)** Correlation plot comparing differential ATAC-seq accessibility to differential H3K27ac CUT&Tag-enriched regions. Axes represent log2 fold changes between skin AA-T_RE_ and sdLN T_MEM_ cells. Each dot represents an individual peak. **(C)** Homer motif analysis on AA-T_RE_ H3K4me1 DER regions. **(D)** Aggregated signal profiles (top) and heatmaps (bottom) showing normalized RUNX2 and RUNX3 CUT&Tag enrichment across consensus peak regions. Consensus peak regions were generated by merging peaks identified across all conditions and were ranked by decreasing mean normalized CUT&Tag signal across the displayed conditions. **(E)** Principal component analysis (PCA) of RUNX2 and RUNX3 CUT&Tag peak counts. Samples are colored by cell population and shaped by TF. **(F)** CUT&Tag tracks showing histone modifications (H3K4me3, H3K27ac, H3K27me3) and TF binding (RUNX2, RUNX3) at the indicated gene loci in indicated cell populations. **(G)** Heatmap representing cumulative CUT&Tag signals associated with the selected genes. For each gene, the signals from all assigned TF-binding peaks were summed to generate a cumulative peak score. **(H)** Bar plot showing the count (left) and percentages (right) of RUNX2- and RUNX3-bound peaks classified as Active (H3K4me1^+^H3K27ac^+^H3K27me3⁻), Primed (H3K4me1^+^H3K27ac⁻ H3K27me3⁻), Repressed (H3K27ac⁻ H3K27me3^+^) chromatin states. DER, differentially enriched region.

Projection onto published T_MEM_ and T_EX_ enhancer programs from chronic LCMV infection (Huang et al., 2025) showed increased representation of T_EX_-associated enhancer states in skin AA-T_RE_ cells relative to sdLN T_MEM_ cells (Fig. S10A-B, Table S1), indicating partial overlap with chronic activation programs. This overlap was selective rather than global. The -23.8 kb *Pdcd1* enhancer, which is associated with T_EX_ but is also accessible in skin T_RM_ cells (Buquicchio et al., 2024), adopted an active configuration in AA-T_RE_ cells (Fig. S10C, left). In contrast, the *Tox* locus remained H3K27me3-enriched and lacked the active enhancer configuration observed in T_EX_ cells (Fig. S10C, middle), consistent with the low TOX expression measured phenotypically.

To define the timing of direct RUNX-family engagement, we performed RUNX2 CUT&Tag in sdLN T_N_, T_MEM_, and AA-T_RE_ cells, as well as in skin AA-T_RE_ cells, and compared these profiles with those of RUNX3. RUNX2 occupancy increased progressively along the trajectory, with substantial binding already present in the sdLN (Fig. 6D). RUNX3 showed moderate genome-wide occupancy in sdLN populations but increased sharply in skin AA-T_RE_ cells (Fig. 6D). PCA confirmed convergence of RUNX2 and RUNX3 binding profiles after differentiation, with the greatest separation in earlier sdLN states (Fig. 6E). At representative effector and residency loci, including *Ifng*, *Itgae*, and *Cd101*, RUNX2 and RUNX3 co-occupied regulatory regions in skin AA-T_RE_ cells (Fig. 6F). Crucially, RUNX2 already occupied T_RE_-associated regulatory loci in the sdLN, whereas RUNX3 was not detectably engaged at these loci until skin entry (Fig. 6F-G). Peak-state analysis similarly showed greater RUNX2-only binding in the sdLN and increased RUNX2/RUNX3 co-occupancy in the skin (Fig. 6H). These data provide direct evidence that RUNX2 engages the AA-T_RE_ program before tissue entry, whereas RUNX3 is recruited to the same regulatory architecture later in the skin. Collectively, histone and TF CUT&Tag analyses support a model in which RUNX2 licenses the resident-effector program in the sdLN before tissue entry, with RUNX3 subsequently reinforcing shared regulatory loci in the skin. Together with the AP-1 motif enrichment and functional perturbation data, these results define a spatially ordered RUNX2- RUNX3-AP-1 regulatory program for AA-T_RE_ differentiation.

### Human AA CD8 T cells share the RUNX2–RUNX3–AP-1 regulatory architecture

Finally, we asked whether key features of the mouse AA-T_RE_ program are present in human AA. We analyzed publicly available scRNA-seq and scATAC-seq datasets from CD8 T cells in AA PBMCs and skin (GSE277469 and GSE212450) (Gay-Mimbrera et al., 2025; Ober-Reynolds et al., 2023). Integrated scRNA-seq identified six clusters representing T_N_, T_CM_, T_EM_, and skin AA-T_RE_-like populations (Fig. 7A-C). Because functional residency and pathogenicity cannot be established from these human datasets, we refer to the skin populations as AA-T_RE_-like. These cells expressed tissue-residency and AP-1-associated genes, including *CD69*, *PRDM1*, *RUNX3*, *FOSL2*, and *JUND*, with low detection yet expression of *CD49a* and *CD103* (Fig. 7C). RUNX2 transcripts were sparse, but we directly detected RUNX2 protein in approximately 60% of perifollicular CD8 T cells in analyzed AA biopsies (n = 3 patients; Fig. 7D). AA-T_RE_-like cells showed little TOX expression, increased TCR-induced activation scores, reduced IL-15-induced activation scores, and increased TGFβ signaling relative to circulating populations (Fig. 7C, E-G). Public scATAC-seq further showed increased accessibility and TF footprints at RUNX2, RUNX3, and FOSL2 motifs in skin AA-T_RE_-like cells (Fig. 7H-J). Thus, human AA CD8 T cells share key transcriptional, protein, and chromatin features of the mouse RUNX2-RUNX3-AP-1 resident-effector program, although a causal requirement for this regulatory axis in human disease remains to be established.

**Figure 7.**
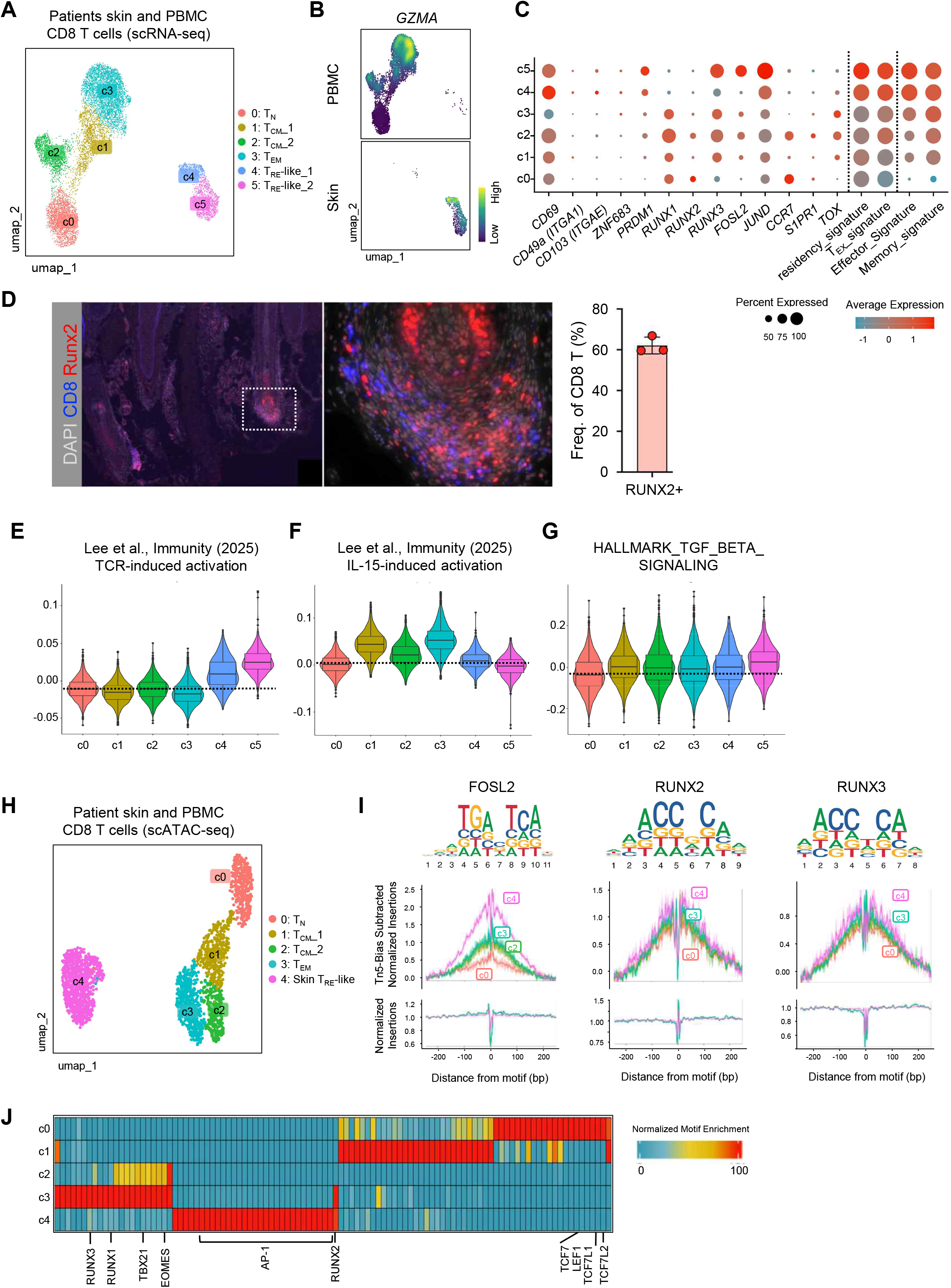
Human AA CD8 T cells share the RUNX2–RUNX3–AP-1 regulatory architecture. **(A-C)** scRNA-seq data of CD8 T cells of the AA patient scalp samples (GSE212450) and PBMCs (GSE277469). UMAP showing indicated CD8 T cells in PBMCs and scalp (A) and density plots showing GZMA expression separated by organ (B). Dot plot of gene expression and pathway enrichment in each cluster. Dot size indicates the fraction of cells expressing the indicated genes; color represents average expression (C). **(D)** Immunofluorescence staining shows the expression pattern of the indicated molecules in the human AA skin (left). Graph shows RUNX2^+^ cells among total CD8 T cells quantified using a histocytometric algorithm (right). **(E-G)** Violin plot showing modular scores of indicated gene sets across AA patient CD8 T cells. **(H-J)** scATAC-seq data of CD8 T cells of AA patient skin samples (GSE212450) and PBMCs (GSE277469). UMAP showing the indicated CD8 T cells in PBMCs and scalp (H). Motif position weight sequence and TF footprinting for the indicated TFs are shown (I). The heatmap shows TF motif enrichment in the marker peaks of the indicated populations (J). PBMC, peripheral blood mononuclear cells

## DISCUSSION

This study identifies a distinct architecture for chronic CD8 T cell-mediated autoimmunity in AA: pathogenicity is maintained by continuous turnover of a terminal resident-effector pool rather than by persistence of the terminal effectors themselves. CD103⁺CD101⁺ AA-T_RE_ cells are tissue resident, clonally selected, TCR-dependent, and short-lived; transferred T_RE_ cells disappear, while host-derived T_RE_ cells accumulate first in the sdLN and subsequently in the skin. Mechanistically, RUNX2 is required for T_RE_ generation and engages the resident-effector regulatory program in the sdLN, whereas RUNX3 recruitment and AP-1 activity become prominent after skin entry. These findings connect the maintenance of chronic disease to repeated production of a terminal pathogenic state and identify the sdLN as the site at which this fate is licensed.

AA-T_RE_ cells also diverge from established skin T_RM_ programs. Skin CD8 residency can be supported by an IFNγ-associated T_RM_1 circuit that depends on T-bet, Hobit, and IL-15, or by an IL-17-associated T_RM_17 circuit linked to ICOS, c-Maf, and IL-7 (Park et al., 2023). AA-T_RE_ cells fit neither pattern in that they are T-bet-high but CD122-low, produce IFNγ rather than IL-17, and further downregulate CD127 during terminal differentiation. This receptor profile is consistent with contraction rather than durable memory, although it should not be interpreted as the direct cause of apoptosis. Notably, *Il2rb* transcripts remain detectable, and RUNX2/RUNX3 occupy the *Il2rb* locus (Fig. 6G) despite loss of CD122 protein (Fig. 3C), suggesting post-transcriptional regulation of IL-15 responsiveness. Defining the mechanism responsible for this discordance may clarify why AA-T_RE_ cells adopt a resident yet short-lived fate.

AA-T_RE_ cells are also distinct from canonical exhausted CD8 T cells. They express multiple inhibitory receptors but retain cytokine production and metabolic activity, show little TOX expression, and maintain the *Tox* locus in an H3K27me3-enriched configuration. RUNX2 overexpression has been reported to suppress TOX and reinforce effector function in antigen-experienced CAR T cells (DeGolier et al., 2025), while sustained AP-1 activity can oppose partnerless NFAT-driven exhaustion (Martinez et al., 2015). These observations are consistent with the RUNX2/AP-1 program identified here. However, TOX was not a direct RUNX2 or RUNX3 binding target in our data, and we did not directly map AP-1 occupancy at the *Tox* locus. Thus, RUNX2 and AP-1 provide a plausible mechanism for maintaining effector competence without canonical exhaustion, but a direct causal link to Tox repression remains to be established.

The short lifespan of AA-T_RE_ cells may also be relevant to the clinical behavior of AA. In contrast to vitiligo, in which long-lived T_RM_ cells can maintain disease for prolonged periods (Frisoli et al., 2020; Richmond et al., 2019; Seneschal et al., 2025), spontaneous recovery is common in AA (Fukuyama et al., 2022). A pathogenic compartment that depends on continuous replenishment rather than durable tissue persistence could provide one explanation for this difference. This architecture also shifts the therapeutic control point: preventing the generation of new AA-T_RE_ cells in the sdLN may be more effective than attempting to eliminate every established skin effector. The reduction in disease incidence after CADD522 treatment provides proof of principle for targeting the differentiation program, although the translational specificity of this approach requires further evaluation.

Our data support a working model in which transferred AA-T_RE_ cells initiate follicular injury but rapidly disappear, after which newly recruited host clonotypes are activated in the sdLN. We speculate that hair-follicle antigens released during tissue injury are presented in a TGFβ-rich draining-node environment, where TCR signaling induces RUNX2 and licenses the T_RE_ program. These cells then enter the skin, where RUNX3 and AP-1 reinforce terminal effector differentiation before the CD39⁺ population undergoes contraction. This model is consistent with the observed kinetics, TCR dependence, and spatial TF engagement, but the initiating autoantigen and the signals that determine the divergence between short-lived AA-T_RE_ cells and durable T_RM_ cells remain unknown. Persistent antigen exposure alone is unlikely to be sufficient, because pmel-1 cells in MAV encounter ongoing autoantigen yet form long-lived T_RM_ populations.

Several limitations define the next steps. First, the identification of the cognate autoantigen(s) will permit direct testing of antigen specificity and tissue homing. Second, although RUNX2 is engaged earlier than RUNX3 and both are individually required for T_RE_ generation, whether RUNX2 acts hierarchically upstream of RUNX3 remains to be determined by genetic epistasis. Third, human AA samples recapitulate key transcriptional, chromatin, and protein features of the mouse program, but the causal requirement of the RUNX2-RUNX3-AP-1 axis in human disease remains to be established through functional perturbation.

Collectively, these findings define AA as a TCR-dependent autoimmune process sustained by the continuous generation of short-lived resident effectors rather than by the persistence of the terminal pathogenic pool. RUNX2 licenses this fate in the sdLN, and RUNX3 together with AP-1 reinforces the resident-effector program after skin entry. Therefore, targeting pathogenic T_RE_ cell production in the sdLN, rather than effector longevity in the skin, would be a control point for therapeutic intervention.

## MATERIALS AND METHODS

### Study design

This study aimed to define the identity, differentiation trajectory, and regulatory mechanisms of AA-T_RE_ cells. Using spontaneous and adoptive-transfer mouse models, we characterized AA-T_RE_ cells by multiplex flow cytometry, parabiosis, adoptive transfer, and functional assays. IBEX imaging, single-cell transcriptomics, TCR repertoire, chromatin accessibility, and CUT&Tag analyses were performed to define the transcriptional and epigenetic programs underlying AA-T_RE_ differentiation. Candidate regulators were validated by CRISPR-Cas9-mediated gene disruption and pharmacologic inhibition. Conservation of the identified differentiation program was assessed using publicly available human single-cell datasets and human AA skin samples.

### Mice

C3H/HeJ mice were purchased from the Jackson Laboratory and maintained under specific-pathogen-free conditions at Seoul National University. B6.CD45.1 (B6-LY5.2/Cr) and B6 *Rag1* KO mice were obtained from Dr. Yeonseok Chung (Seoul National University), and B6 T-bet reporter mice with ZsGreen were described previously.(Lee et al., 2013) CD45.1, *Rag1* KO, and T-bet^ZsGreen^ mice in a B6 background were backcrossed to the C3H/HeJ mice for at least 10 generations before use. All mouse experiments were performed in accordance with protocols approved by the Institutional Animal Care and Use Committee (IACUC) of Seoul National University.

### Human samples

Patients with alopecia areata who underwent punch biopsy at Seoul National University Hospital between January 2023 and March 2026 were retrospectively enrolled. The study protocol was approved by the Institutional Review Board of Seoul National University Hospital (IRB No. H-2604-217-1745), which granted a waiver of informed consent for the use of archived FFPE specimens, and the study was conducted in accordance with the Declaration of Helsinki. Formalin-fixed, paraffin-embedded (FFPE) tissue blocks from eight patients were sectioned at a thickness of 4 μm. Immunohistochemical staining was performed using anti-RUNX2 (CST#12556) and anti-CD8 (BioLegend #372908) antibodies with the SuperBoost kit (Invitrogen#B40916, Invitrogen#B40923). Three samples containing abundant CD8 T cells were analyzed. Publicly available human scRNA-seq (GSE277469) and scATAC-seq (GSE212450) datasets were reanalyzed as described in Fig. 6 under their respective institutional approvals.

### C3H/HeJ model of AA

AA mice were generated as previously described.(Lee et al., 2022) Briefly, sdLNs were removed from AA mice, and cells were cultured with AR10 (advanced RPMI 1640, Gibco) containing 10% FBS (Atlas Biologicals), 2mM GlutaMAX (Gibco), and 100 U/mL Penicillin-Streptomycin-Tetracycline (Welgene) supplemented with rmIL-2, rmIL-7, and rmIL-15 cytokines (Genscript). Cells were stimulated with anti-CD3 and anti-CD28-coated microbeads (Dynabeads, Thermo Fisher) for 7 days and intradermally transferred into at least 10-week-old normal-haired female C3H/HeJ mice. In some experiments, CD8 T cells isolated from the skin and sdLN were injected intradermally into UA hosts, yielding a similar AA incidence.

### Flow cytometry

Single-cell suspensions were prepared from thymi, spleens, and sdLNs. Mononuclear cells from the liver and lungs were separated using Percoll gradients as described previously.(Yang et al., 2021) Foxp3 staining buffer set (eBioscience) was used to stain intracellular TFs and cytokines. To detect apoptotic cells, Annexin V Apoptosis Detection Kit (Thermo Fisher) was used, and all tissues were prepared using identical enzymatic digestion. Cells were analyzed on an LSR II (Becton Dickinson) or Cytek Aurora 5 (Cytek Biosciences), and data were processed with FlowJo software (TreStar).

### Parabiosis

Age- and sex-matched UA- or AA–induced C3H/HeJ mice with ≤5% body weight difference were paired. Mice were anesthetized by intraperitoneal injection of ketamine and xylazine (100 mg/kg and 10 mg/kg, 10 µL/g body weight). An ocular ointment was applied, and body temperature was maintained with a heating pad. The lateral aspects from the axilla to the mid-thigh were shaved, and corresponding skin incisions were aligned and sutured using 4-0 nonabsorbable nylon sutures and surgical clips (BD). Postoperative analgesia was provided with buprenorphine (0.05 mg/kg, subcutaneously, every 12 h) for 48–72 h, and mice were monitored at least twice daily for the first 3 days. Successful circulatory sharing was confirmed 2–3 weeks after surgery by flow cytometric detection of partner-specific CD45 allotypes in peripheral blood leukocytes.

### BrdU assay

Mice were injected intraperitoneally with 1mg BrdU 16 h before tissue harvest. BrdU incorporation was detected using the BrdU Staining Kit (Invitrogen #8812-6600) according to the manufacturer’s instructions, following surface marker staining.

### In vitro CD8 T cell differentiation

CD8 T_MEM_ cells from sdLN of unaffected C3H/HeJ mice were sorted and then cultured for 3 days in the presence of the indicated cytokine combination. IL-12 (50ug/ml, R&D systems), IL-15 (50ng/ml; GenScript), IL-18 (50ug/ml, R&D systems), TGFβ (5ng/ml, Peprotech), and Mouse T-Activator CD3/28 Dynabeads (ThermoFisher) were used.

### LCMV infection

For chronic LCMV infection, mice were infected with 2 × 10^6^ plaque-forming units of LCMV clone 13 (Cl-13) intravenously. Transient CD4 T cell depletion was performed at the onset of infection by injecting 200 μg of anti-CD4 depleting antibodies (GK1.5, BioXCell) intraperitoneally 2 days before and on the day of infection.

### CRISPR-Cas9 editing and cell transfer

CRISPR-Cas9 editing. crRNA targeting: Thy1 (5’-CGTGTGCTCGGGTATCCCAA-3’), NKG2D (5’-GCACTAACTACCAGTCAACC-3’, 5’-AAGGGTGAATCGAATTGCCA-3’), TRAC (5’-TTCGGAGTCCCATAACTGAC-3’), TRBC (5’-GTTGCGAGGATTGTGCCAGA-3’) and tracrRNA were purchased from Integrated DNA Technologies (IDT). Single-guide RNA duplexes (sgRNAs, crRNA-tracrRNA duplexes) were formed by incubating 0.3 nmol each of crRNA and tracrRNA at 95℃ for 5 minutes, then cooling to RT for 1 hour. RNPs were formed by adding 1.62µl Alt-R S.p. Cas9 Nuclease V3 (10mg/ml; IDT) to the sgRNAs (molar ratio of 3:1= sgRNA: Cas9 nuclease) and incubating for 10 minutes at RT. CD8 T cells were activated in vitro with Mouse T-Activator CD3/28 Dynabeads (Thermo Fisher) for 48 hours. After removing the beads, the in vitro-activated CD8 T cells were resuspended in 20 µL of P3 solution (P3 Primary Cell 4D-Nucleofector X Kit S; Lonza) and incubated with the RNP complex for 2 minutes at RT. The mixture was electroporated using the Lonza 4D-Nucleofector system (DN100). Cells were expanded for an additional 4 days in the presence of IL-2 (50 U/ml; National Institutes of Health), IL-7 (25 ng/ml; GenScript), and IL-15 (50 ng/ml; GenScript). Seven to ten million edited cells were injected intradermally into C3H recipients.

### In vivo Runx2 blocker treatment

UA C3H/HeJ mice were injected with a Runx2 blocker (CADD522, 100 mg/kg, MedChemExpress) or vehicle (corn oil, Sigma-Aldrich) once weekly for 5 weeks, starting at the time of induction. To test the effect, the mice were treated once weekly for 5 consecutive weeks.

### Oxygen Consumption Rate (OCR)

One day before the assay, a Seahorse XFe96 Cell Culture Microplate (Agilent) was coated with Poly-D-Lysine (50 µg/mL) overnight at 4°C, and the Sensor Cartridge (Agilent) was hydrated with XF Calibrant (Agilent) overnight at 37°C. On the day of the actual assay, T_N_, T_MEM_, and AA-T_RE_ CD8 T cells were sorted from AA mice and seeded into a Cell Culture Microplate. 2∼3 x 10^5^ T cells were used per well, and at least 4 replicates were made. Drug compounds were prepared according to the manufacturer’s protocols, and analysis was performed using an Agilent Seahorse XFe96 Analyzer.

### IBEX imaging

IBEX imaging was performed as previously described.(Radtke et al., 2022) Briefly, mouse tissues were fixed in 4% PFA (EMS) for 30 min, incubated overnight in 30% sucrose, and snap-frozen. Two-well chambered cover glasses (Thermo Fisher Scientific) were coated with 15 μL of chrome alum gelatin (Newcomer Supply). The coated cover glasses were dried for 1 hr at 60 °C in the oven. Tissue sections were cut directly onto the coated cover glasses and dried for 1 hr at 37 °C. Sections were rehydrated by adding 1 mL phosphate-buffered saline (PBS) per well for 5 minutes. Tissue sections were blocked with 5% bovine serum albumin and goat and mouse sera (Jackson Laboratory) for 1 hr at 25 °C. Primary antibody solution was added and incubated for 1 hr at 37 °C. Samples were washed again with PBS and mounted in 1 mL of Fluoromount-G (SouthernBiotech). After imaging, Fluoromount-G was removed by suction, followed by three PBS exchanges. For fluorochrome bleaching, lithium borohydride (LiBH₄; Strem Chemicals) solution was added to the tissue, which was then incubated for 10 min at 25 °C. The tissue was washed three times with PBS. The imaging steps were repeated until the desired number of antibody panels was achieved.

### Histocytometry

Histocytometric analysis was performed as described previously.(Lee et al., 2015) Briefly, the fluorochrome intensities for each region of interest were quantified using ImageJ, and the data were exported to Excel and Prism for two-dimensional plotting. XY coordinates of gated populations were overlaid on original images to analyze localization.

### Peripheral T cell retroviral transduction

Single-cell suspensions from sdLNs and spleen were enriched for CD8 T cells using anti-CD8 microbeads (Miltenyi Biotec). Purified CD8 T cells were resuspended in T cell culture medium, and 1 × 10⁶ cells were plated per well in a 24-well cell culture plate (VWR), supplemented with 100 U/mL recombinant human IL-2 (hIL-2; NIH) and Dynabeads Mouse T-Activator CD3/CD28 (Gibco). Cells were incubated at 37 °C overnight for activation. Activated CD8 T cells were transduced with a retrovirus encoding the TCR of interest and a GFP reporter in the presence of hIL-2 and 10 µg/mL polybrene (Sigma-Aldrich). Spin infection was performed at 2,000 × g for 90 min at 37 °C after Dynabeads were removed. Following transduction, 1 mL of fresh T cell culture medium containing hIL-2 was added immediately, and cells were incubated for 24 h. Cells were then centrifuged, resuspended in T cell culture medium with hIL-2, and cultured for an additional 48 h. Transduced CD8⁺ T cells were washed once with PBS, resuspended in PBS, and 5–10 × 10⁶ cells were injected intravenously or intradermally into recipient C3H/HeJ mice.

### Melanoma-associated vitiligo (MAV) model

MAV was generated as previously described.(Molodtsov et al., 2021) Briefly, 1 x 10^4^ naïve pmel-1 CD8 T cells were adoptively transferred intravenously into wild-type C57BL/6 mice. One day later, mice were inoculated intradermally with 1 x 10^5^ B16-F10 melanoma cells. Recipient mice received 250 µg anti-CD4 mAb on days 4 and 10, and primary tumors were surgically resected on day 12. Mice that subsequently developed depigmentation were used as the MAV group for downstream analyses.

### Cicatricial alopecia model

To generate TCR retrogenic mice, TCR⍺ (TRAV8D-2/TRAJ37) and TCRβ (TRBV13-2/TRBJ2-7) chains were cloned into a pMSCV-IRES-GFP retroviral backbone.(Alli et al., 2012) Platinum-E cells were transfected with the construct using FuGENE HD (3:1 ratio), and the resulting viral supernatant was concentrated overnight with RetroX Concentrator (1:3 v/v) and resuspended in BM media. For donor cell preparation, 8–9-week-old female C57BL/6 *Rag1*^-/-^CD4^Cre^ mice received an intraperitoneal injection of 5- fluorouracil (0.15 mg/g). After 72 hours, BM cells were flushed from femurs and tibiae and cultured in BM media supplemented with rmIL-3 (20 ng/mL), rhIL-6 (50 ng/mL), and rmSCF (50 ng/mL). These cells were transduced with the TCR-expressing retrovirus via two consecutive spinoculations (1000 x g for 1 hour at 37 °C) with polybrene (10 µg/mL) over 48 hours. Finally, transduced BM cells were retro-orbitally injected into sublethally irradiated *Rag1*^-/-^recipients, which were monitored for 10 weeks for hair loss and general health.

### 10x Genomics library generation and sequencing

Sequencing libraries for scRNA-seq and scTCR-seq were generated using Chromium Next GEM Single Cell 5’ Reagent Kits (10x Genomics). Three libraries were generated for each sample using the 5’ RNA library kit, 5’ RNA Feature Barcode for HTO kit, and Single Cell VDJ library v2 (10x Genomics). Sequencing libraries for multiome sequencing were generated using Chromium Next GEM Single Cell Multiome ATAC^+^ Gene Expression Reagent (10X Genomics).

### Single-cell RNA and TCR sequencing

Alignment, barcode counting, and feature-barcode matrices were generated using Cell Ranger 7.0.1. Data were further analyzed using Seurat (v5.1.0).(Hao et al., 2024) All single-cell analysis was performed using R (v4.4.1). Raw count data were QC-filtered according to a protocol developed by the Harvard Chan Bioinformatics Core (HBC).(Mary Piper, 2022) In brief, cells were filtered according to RNA counts, gene counts, log10GenesPerUMI, and mitochondrial gene percentage. Genes expressed in fewer than 10 cells were removed, along with TCR genes, to reduce bias in clustering. Samples from different organs were demultiplexed using deMULTIplex2 (v1.0.2), and doublets were identified using DoubletFinder (v2.0.4).(McGinnis et al., 2019) Samples were merged using Harmony (v1.2.3)(Korsunsky et al., 2019), and cells without TCR clone information were removed for further analysis. V(D)J transcripts were assembled using Cell Ranger v 7.0.1. and further analyzed in R using scRepertoire (v2.2.1).(Yang et al., 2025) All TCR analyses were conducted on cells with available gene expression information.

### Single-cell multiome analysis

Alignment, barcode counting, and feature-barcode matrices were generated using Cell Ranger 8.0.1. Data were further analyzed using Seurat (v.5.1.0) and Signac (v1.14.0).(Hao et al., 2024; Stuart et al., 2021) Seurat objects were created from raw count data using Seurat (v.5.1.0). The ATAC data were added to the Seurat object using CreateChromatinAssay (Seurat). Quality control for the combined data was performed by filtering cells based on transcription start site enrichment scores, nucleosome signal scores, the percentile of reads in peaks, total ATAC counts, total RNA counts, total gene counts, the percentile of mitochondrial counts, and log10GenesPerUMI. Integrated UMAP embeddings that represent gene expression and chromatin accessibility profiles were constructed using FindMultiModalNeighbors (Seurat), which implements the weighted nearest neighbor (WNN) method to capture multimodal relationships among cells. Motif enrichment analysis was performed using chromVAR v.1.28.0, using JASPAR2026.(Ovek Baydar et al., 2025; Schep et al., 2017)

### SCENIC_+_ analysis

Integrated single-cell transcriptomic and single-cell chromatin accessibility data from the Seurat analysis were used to identify gene regulatory networks using SCENIC^+^ (v1.0a2).(Bravo Gonzalez-Blas et al., 2023) Genome annotation and chromosome size files were retrieved using biomaRt (http://ensembl.org/biomart) with the Mus musculus reference. eRegulon results were filtered to retain only those with a correlation > 0.7 between gene-based and region-based regulon area under the curve values. The resulting gene regulatory networks were visualized using Cytoscape.(Shannon et al., 2003)

### ArchR analysis

ATAC-seq fragment files for each sample from public repositories were processed using ArchR (v1.0.2).(Granja et al., 2021) Quality control of patient scalp data was performed using the values provided in the original paper. Quality control for multiome data from patient peripheral blood was performed by filtering cells based on transcription start site enrichment scores, nucleosome signal scores, percentile of reads in peaks, total ATAC counts, total RNA counts, total gene counts, percentile of mitochondrial counts, and log10GenesPerUMI. CD8 T cells identified by marker features were selected for downstream analysis. Motif enrichment analysis in differential peaks was performed using JASPAR2020 (v0.99.8).

### ATAC-seq

Bulk ATAC-seq was performed on sorted cells as described (Grandi et al., 2022) with some optimizations. Briefly, sorted cells were adjusted to approximately 1 × 10⁶ cells/mL, and 50,000 cells were used per reaction. To control technical variation across samples, Drosophila S2 cell nuclei were added as an internal spike-in control (10,000 nuclei per reaction) at the initial lysis step. Nuclei were isolated under detergent conditions and subjected to transposition using Tn5 transposase (Enzynomics). Following tagmentation, DNA was purified using MinElute PCR Purification Columns (QIAGEN) and amplified with indexed primers. Library amplification cycles were empirically determined by qPCR to minimize overamplification. Libraries were purified using MinElute PCR purification columns (QIAGEN) and assessed for fragment size distribution and concentrations before sequencing.

### CUT&Tag-seq library construction

CUT&Tag-seq was performed on sorted cells as described (Kaya-Okur et al., 2019), with some modifications based on the EpiCypher CUTANA™ and Vazyme Hyperactive® Universal CUT&Tag Assay Kit for Illumina Pro CUT&Tag platforms, following the manufacturers’ instructions. For histone modification profiling, cells were lightly cross-linked with 0.1% PFA for 1 minute at room temperature, and nuclei were isolated. For each reaction, 100,000 nuclei were bound to Concanavalin A magnetic beads (Epicypher, 21-1401). Primary antibodies against histone modifications (H3K4me1, CST#5326, 1:100; H3K4me3, Abcam # ab8580, 1:100; H3K27ac, Active Motif# 39133, 1:100; H3K27me3, CST#9733, 1:50) were incubated with bead-bound nuclei overnight at 4°C with agitation, followed by secondary antibody binding the next day. For transcription factor profiling, 200,000 sorted cells were processed using the Hyperactive Universal CUT&Tag Assay Kit for Illumina Pro (Vazyme, TD904-01). Cells were subjected to nuclei isolation and light cross-linking using the Hyperactive TF Enhancer Module for CUT&Tag (Vazyme, TD904-C2). Isolated nuclei were bound to Concanavalin A magnetic beads and incubated overnight at 4°C with 1 µg primary antibodies against TFs (RUNX2, CST#12556; RUNX3, Invitrogen#MA5-17169; Rabbit IgG Isotype Control, Invitrogen#02-6102), followed by secondary antibody binding the next day. Following secondary antibody binding, Protein A/G–Tn5 transposase loaded with sequencing adapters was then added to tether Tn5 to antibody-bound chromatin. Tagmentation was initiated under Mg²⁺-containing conditions, and DNA fragments were released. CUT&Tag libraries were PCR-amplified, purified using AMPure XP beads, and assessed for fragment size distribution and concentration before sequencing on an Illumina platform.

### ATAC-seq and CUT&Tag-seq data analysis

Libraries were sequenced in 150 bp paired-end mode on an Illumina NovaX platform. Raw sequencing reads were adapter-trimmed using Trim Galore in paired-end mode and subsequently aligned to the mouse reference genome (mm10) using bowtie2 (paired-end; maximum fragment length 1000 bp; one mismatch seed setting -N 1) (Langmead and Salzberg, 2012). Aligned reads were converted to BAM format, sorted, and indexed using samtools (Li et al., 2009), and PCR duplicates were removed using Picard. Uniquely and appropriately paired reads (MAPQ ≥ 20) were retained for downstream analyses. Genome-wide signal tracks were generated from filtered BAM files as bigWig files using deepTools with 10 bp binning, applying RPGC normalization (mm10 effective genome size = 2.65 × 10⁹). Peak calling was performed using MACS3 in paired-end mode (Zhang et al., 2008). Histone marks with focal enrichment (e.g., H3K27ac, H3K4me3, ATAC) were called as narrow peaks, whereas broader marks (e.g., H3K4me1, H3K27me3) were called using broad peak settings. Downstream analyses were performed using R (v4.5.1), including differential accessibility and enrichment analysis with DiffBind/DESeq2, peak overlap and annotation using GenomicRanges, and customized visualization workflows integrating deepTools (v3.5.6) outputs to compare chromatin accessibility and histone modification dynamics across conditions.(Ramirez et al., 2016)

### Statistical analyses

All experimental data plotting and statistical analyses were performed using GraphPad Prism (v11.0.2). Statistical methodologies are explicitly described in the corresponding figure legends.

## Supplementary Materials

Fig. S1 - S11.

Table S1.

## Acknowledgments

This work was supported by the Samsung Science & Technology Foundation (SSTF-BA2202-08).

## Author contributions

YL and JC designed and performed experiments and analyzed data; MSL performed TCR cloning and retroviral transduction; JC, SK, JJK and JYL performed CUT&Tag analysis; SP, and SJ analyzed RNA-seq data; SJS generated B6 retrogenic mice; THK performed immunofluorescence staining; SHJ performed mouse experiments and data analysis; KL and SJI performed LCMV infection; CL, HL and OK provided human samples; SJI and JYL provided research interpretation; JYL and YJL wrote the manuscript; YJL conceptualized and supervised the research.

## Competing interests

The authors declare no competing financial interests.

## Data and materials availability

Mice and reagents used in this study are available upon request. Raw and processed sequencing data from RNA-seq, ATAC-seq, and CUT&Tag-seq have been deposited in the National Center for Biotechnology Information Gene Expression Omnibus under accession codes GSE320508 and GSE322667. Publicly available datasets analyzed during this manuscript were GSE233389, GSE151652, GSE212450, GSE277469, or via https://data.mendeley.com/datasets/gxhhxrtr2w/1. Source data are provided with this paper. Requests for additional information and resource requests will be fulfilled by the lead contact, You Jeong Lee.

## Notes

### Competing Interest Statement

The authors have declared no competing interest.

## REFERENCES

Al Souz, J., Y. Wei, C. Cui, R. Qu, J.Y. Choi, F. Wang, S.J. Moioffer, J. Barrett, I. Chernova, M. Nadella, E.R. Siniscalco, G. Moeckel, N.S. Joshi, P.M. Chen, A. Arazi, R.A.S.L.E.N. Accelerating Medicines Partnership, Y.C. Ho, and J. Craft. 2026. Organ injury in systemic autoimmunity is mediated by stem-like CD8(+) T cells arising from tissue-draining lymph nodes. Immunity 59:1667–1687 e1615.

Alli, R., P. Nguyen, K. Boyd, J.P. Sundberg, and T.L. Geiger. 2012. A mouse model of clonal CD8+ T lymphocyte-mediated alopecia areata progressing to alopecia universalis. J Immunol 188:477–486.

Belk, J.A., B. Daniel, and A.T. Satpathy. 2022. Epigenetic regulation of T cell exhaustion. Nat Immunol 23:848–860.

Bravo Gonzalez-Blas, C., S. De Winter, G. Hulselmans, N. Hecker, I. Matetovici, V. Christiaens, S. Poovathingal, J. Wouters, S. Aibar, and S. Aerts. 2023. SCENIC+: single-cell multiomic inference of enhancers and gene regulatory networks. Nat Methods 20:1355–1367.

Buquicchio, F.A., R. Fonseca, P.K. Yan, F. Wang, M. Evrard, A. Obers, J.C. Gutierrez, C.J. Raposo, J.A. Belk, B. Daniel, P. Zareie, K.E. Yost, Y. Qi, Y. Yin, K.F. Nico, F.M. Tierney, M.R. Howitt, C.A. Lareau, A.T. Satpathy, and L.K. Mackay. 2024. Distinct epigenomic landscapes underlie tissue-specific memory T cell differentiation. Immunity

Byrne, K.T., A.L. Cote, P. Zhang, S.M. Steinberg, Y. Guo, R. Allie, W. Zhang, M.S. Ernstoff, E.J. Usherwood, and M.J. Turk. 2011. Autoimmune melanocyte destruction is required for robust CD8+ memory T cell responses to mouse melanoma. J Clin Invest 121:1797–1809.

Dai, Z., Y. Chang, E.Y. Lee, V.P. Leifer, E.H. Chun Wang, R. Perez-Lorenzo, A. de Jong, and A.M. Christiano. 2025. Single-cell analysis of temporal immune cell dynamics in alopecia areata reveals a causal role for clonally expanded CD8(+) T cells in disease. Cell Rep 44:115798.

de Jong, A., A. Jabbari, Z. Dai, L. Xing, D. Lee, M.M. Li, M. Duvic, M. Hordinsky, D.A. Norris, V. Price, J. Mackay-Wiggan, R. Clynes, and A.M. Christiano. 2018. High-throughput T cell receptor sequencing identifies clonally expanded CD8+ T cell populations in alopecia areata. JCI Insight 3:

DeGolier, K.R., E. Danis, M. D’Antonio, J. Cimons, M. Yarnell, R.M. Kedl, M.E. Kohler, J.P. Scott-Browne, and T.J. Fry. 2025. Antigen experience history directs distinct functional states of CD8(+) CAR T cells during the antileukemia response. Nat Immunol 26:68–81.

Dillon, K.L. 2021. A Comprehensive Literature Review of JAK Inhibitors in Treatment of Alopecia Areata. Clin Cosmet Investig Dermatol 14:691–714.

Frisoli, M.L., K. Essien, and J.E. Harris. 2020. Vitiligo: Mechanisms of Pathogenesis and Treatment. Annu Rev Immunol 38:621–648.

Fukuyama, M., T. Ito, and M. Ohyama. 2022. Alopecia areata: Current understanding of the pathophysiology and update on therapeutic approaches, featuring the Japanese Dermatological Association guidelines. J Dermatol 49:19–36.

Gavil, N.V., M.C. Scott, E. Weyu, O.C. Smith, S.D. O’Flanagan, S. Wijeyesinghe, S. Lotfi-Emran, S.L. Shiao, V. Vezys, and D. Masopust. 2023. Chronic antigen in solid tumors drives a distinct program of T cell residence. Sci Immunol 8:eadd5976.

Gay-Mimbrera, J., P.J. Gómez-Arias, P. Álvarez-Heredia, A. Batista-Duharte, I. Rivera-Ruiz, M. Aguilar-Luque, M. Juan-Cencerrado, C. Mochón-Jiménez, Á. Cebrián-García, E. Andújar-Pulido, M. Pérez-Alegre, A. Pera, and J. Ruano. 2025. Integrated single-cell chromatin and transcriptomic analyses of peripheral immune cells in patients with alopecia areata. Frontiers in Immunology 16:

Gearty, S.V., F. Dundar, P. Zumbo, G. Espinosa-Carrasco, M. Shakiba, F.J. Sanchez-Rivera, N.D. Socci, P. Trivedi, S.W. Lowe, P. Lauer, N. Mohibullah, A. Viale, T.P. DiLorenzo, D. Betel, and A. Schietinger. 2022. An autoimmune stem-like CD8 T cell population drives type 1 diabetes. Nature 602:156–161.

Gilhar, A., A. Etzioni, and R. Paus. 2012. Alopecia areata. N Engl J Med 366:1515–1525.

Grandi, F.C., H. Modi, L. Kampman, and M.R. Corces. 2022. Chromatin accessibility profiling by ATAC-seq. Nat Protoc 17:1518–1552.

Granja, J.M., M.R. Corces, S.E. Pierce, S.T. Bagdatli, H. Choudhry, H.Y. Chang, and W.J. Greenleaf. 2021. ArchR is a scalable software package for integrative single-cell chromatin accessibility analysis. Nat Genet 53:403–411.

Hao, Y., T. Stuart, M.H. Kowalski, S. Choudhary, P. Hoffman, A. Hartman, A. Srivastava, G. Molla, S. Madad, C. Fernandez-Granda, and R. Satija. 2024. Dictionary learning for integrative, multimodal and scalable single-cell analysis. Nat Biotechnol 42:293–304.

Huang, H., A.E. Baxter, Z. Zhang, C.R. Good, K.A. Alexander, Z. Chen, P.A.A. Garcia, P. Samareh, S.M. Collins, K.M. Glastad, L. Wang, G. Donahue, S. Manne, J.R. Giles, J. Shi, S.L. Berger, and E.J. Wherry. 2025. Deciphering the role of histone modifications in memory and exhausted CD8 T cells. Sci Rep 15:17359.

Kaya-Okur, H.S., S.J. Wu, C.A. Codomo, E.S. Pledger, T.D. Bryson, J.G. Henikoff, K. Ahmad, and S. Henikoff. 2019. CUT&Tag for efficient epigenomic profiling of small samples and single cells. Nat Commun 10:1930.

Khan, O., J.R. Giles, S. McDonald, S. Manne, S.F. Ngiow, K.P. Patel, M.T. Werner, A.C. Huang, K.A. Alexander, J.E. Wu, J. Attanasio, P. Yan, S.M. George, B. Bengsch, R.P. Staupe, G. Donahue, W. Xu, R.K. Amaravadi, X. Xu, G.C. Karakousis, T.C. Mitchell, L.M. Schuchter, J. Kaye, S.L. Berger, and E.J. Wherry. 2019. TOX transcriptionally and epigenetically programs CD8(+) T cell exhaustion. Nature 571:211–218.

Korsunsky, I., N. Millard, J. Fan, K. Slowikowski, F. Zhang, K. Wei, Y. Baglaenko, M. Brenner, P.R. Loh, and S. Raychaudhuri. 2019. Fast, sensitive and accurate integration of single-cell data with Harmony. Nat Methods 16:1289–1296.

Langmead, B., and S.L. Salzberg. 2012. Fast gapped-read alignment with Bowtie 2. Nat Methods 9:357–359.

Lee, E., M. Kim, and Y.J. Lee. 2022. Selective Expansion of Tregs Using the IL-2 Cytokine Antibody Complex Does Not Reverse Established Alopecia Areata in C3H/HeJ Mice. Front Immunol 13:874778.

Lee, E.Y., Z. Dai, A. Jaiswal, E.H.C. Wang, N. Anandasabapathy, and A.M. Christiano. 2023. Functional interrogation of lymphocyte subsets in alopecia areata using single-cell RNA sequencing. Proc Natl Acad Sci U S A 120:e2305764120.

Lee, Y.J., K.L. Holzapfel, J. Zhu, S.C. Jameson, and K.A. Hogquist. 2013. Steady-state production of IL-4 modulates immunity in mouse strains and is determined by lineage diversity of iNKT cells. Nat Immunol 14:1146–1154.

Lee, Y.J., H. Wang, G.J. Starrett, V. Phuong, S.C. Jameson, and K.A. Hogquist. 2015. Tissue-Specific Distribution of iNKT Cells Impacts Their Cytokine Response. Immunity 43:566–578.

Li, H., B. Handsaker, A. Wysoker, T. Fennell, J. Ruan, N. Homer, G. Marth, G. Abecasis, R. Durbin, and S. Genome Project Data Processing. 2009. The Sequence Alignment/Map format and SAMtools. Bioinformatics 25:2078–2079.

Mackay, L.K., A. Rahimpour, J.Z. Ma, N. Collins, A.T. Stock, M.L. Hafon, J. Vega-Ramos, P. Lauzurica, S.N. Mueller, T. Stefanovic, D.C. Tscharke, W.R. Heath, M. Inouye, F.R. Carbone, and T. Gebhardt. 2013. The developmental pathway for CD103(+)CD8+ tissue-resident memory T cells of skin. Nat Immunol 14:1294–1301.

Martinez, G.J., R.M. Pereira, T. Aijo, E.Y. Kim, F. Marangoni, M.E. Pipkin, S. Togher, V. Heissmeyer, Y.C. Zhang, S. Crotty, E.D. Lamperti, K.M. Ansel, T.R. Mempel, H. Lahdesmaki, P.G. Hogan, and A. Rao. 2015. The transcription factor NFAT promotes exhaustion of activated CD8(+) T cells. Immunity 42:265–278.

Mary Piper, M.M., Jihe Liu, William Gammerdinger, Radhika Khetani. 2022. hbctraining/scRNA-seq_online: scRNA-seq Lessons from HCBC (first release). In.

McGinnis, C.S., L.M. Murrow, and Z.J. Gartner. 2019. DoubletFinder: Doublet Detection in Single-Cell RNA Sequencing Data Using Artificial Nearest Neighbors. Cell Syst 8:329–337 e324.

Molodtsov, A.K., N. Khatwani, J.L. Vella, K.A. Lewis, Y. Zhao, J. Han, D.E. Sullivan, T.G. Searles, N.K. Preiss, T.B. Shabaneh, P. Zhang, A.R. Hawkes, B.T. Malik, F.W.t. Kolling, E.J. Usherwood, S.L. Wong, J.D. Phillips, K. Shirai, C.V. Angeles, S. Yan, T.J. Curiel, Y.H. Huang, C. Cheng, and M.J. Turk. 2021. Resident memory CD8(+) T cells in regional lymph nodes mediate immunity to metastatic melanoma. Immunity 54:2117–2132 e2117.

Moran, A.E., K.L. Holzapfel, Y. Xing, N.R. Cunningham, J.S. Maltzman, J. Punt, and K.A. Hogquist. 2011. T cell receptor signal strength in Treg and iNKT cell development demonstrated by a novel fluorescent reporter mouse. J Exp Med 208:1279–1289.

Ober-Reynolds, B., C. Wang, J.M. Ko, E.J. Rios, S.Z. Aasi, M.M. Davis, A.E. Oro, and W.J. Greenleaf. 2023. Integrated single-cell chromatin and transcriptomic analyses of human scalp identify gene-regulatory programs and critical cell types for hair and skin diseases. Nat Genet 55:1288–1300.

Ovek Baydar, D., I. Rauluseviciute, D.R. Aronsen, R. Blanc-Mathieu, I. Bonthuis, H. de Beukelaer, K. Ferenc, A. Jegou, V. Kumar, R.B. Lemma, J. Lucas, M. Pochon, C.M. Yun, V. Ramalingam, S.S. Deshpande, A. Patel, G.K. Marinov, A.T. Wang, A. Aguirre, J.A. Castro-Mondragon, D. Baranasic, J. Cheneby, S. Gundersen, M. Johansen, A. Khan, M.L. Kuijjer, E. Hovig, B. Lenhard, A. Sandelin, K. Vandepoele, W.W. Wasserman, F. Parcy, A. Kundaje, and A. Mathelier. 2025. JASPAR 2026: expansion of transcription factor binding profiles and integration of deep learning models. Nucleic Acids Res

Park, S.L., S.N. Christo, A.C. Wells, L.C. Gandolfo, A. Zaid, Y.O. Alexandre, T.N. Burn, J. Schroder, N. Collins, S.J. Han, S.M. Guillaume, M. Evrard, C. Castellucci, B. Davies, M. Osman, A. Obers, K.M. McDonald, H. Wang, S.N. Mueller, G. Kannourakis, S.P. Berzins, L.A. Mielke, F.R. Carbone, A. Kallies, T.P. Speed, Y. Belkaid, and L.K. Mackay. 2023. Divergent molecular networks program functionally distinct CD8(+) skin-resident memory T cells. Science 382:1073–1079.

Park, S.L., M.M. Painter, S. Manne, V. Alcalde, M. McLaughlin, M.A. Sullivan, D. Mathew, L. Torres, Y.J. Huang, D.B. Reeg, N.R. Douek, T. Campos, M. Klapholz, M.A. Cardenas, V. Fang, S.F. Ngiow, W. Kc, R.R. Goel, A.E. Baxter, J.E. Wu, M. Tan, C.T. Berry, C.T. Ellebrecht, H.C. Alexander, E. Papazian, Y. Liu, K. Rajasekaran, R.M. Brody, E.R. Thaler, D. Basu, A. Diab, J.R. Giles, and E.J. Wherry. 2026. Tissue-resident exhausted and memory CD8(+) T cells have distinct ontogeny, function and role in disease. Nat Immunol 27:110–125.

Pratt, C.H., L.E. King, Jr., A.G. Messenger, A.M. Christiano, and J.P. Sundberg. 2017. Alopecia areata. Nat Rev Dis Primers 3:17011.

Radtke, A.J., C.J. Chu, Z. Yaniv, L. Yao, J. Marr, R.T. Beuschel, H. Ichise, A. Gola, J. Kabat, B. Lowekamp, E. Speranza, J. Croteau, N. Thakur, D. Jonigk, J.L. Davis, J.M. Hernandez, and R.N. Germain. 2022. IBEX: an iterative immunolabeling and chemical bleaching method for high-content imaging of diverse tissues. Nat Protoc 17:378–401.

Ramirez, F., D.P. Ryan, B. Gruning, V. Bhardwaj, F. Kilpert, A.S. Richter, S. Heyne, F. Dundar, and T. Manke. 2016. deepTools2: a next generation web server for deep-sequencing data analysis. Nucleic Acids Res 44:W160–165.

Richmond, J.M., J.P. Strassner, M. Rashighi, P. Agarwal, M. Garg, K.I. Essien, L.S. Pell, and J.E. Harris. 2019. Resident Memory and Recirculating Memory T Cells Cooperate to Maintain Disease in a Mouse Model of Vitiligo. J Invest Dermatol 139:769–778.

Schep, A.N., B. Wu, J.D. Buenrostro, and W.J. Greenleaf. 2017. chromVAR: inferring transcription-factor-associated accessibility from single-cell epigenomic data. Nat Methods 14:975–978.

Seneschal, J., J.M. Bae, K. Ezzedine, I. Hamzavi, J.E. Harris, B. Bellei, D. Parsad, T. Passeron, N. van Geel, K. Boniface, and M. Picardo. 2025. Vitiligo. Nat Rev Dis Primers 11:85.

Seok, J., S.D. Cho, J. Lee, Y. Choi, S.Y. Kim, S.M. Lee, S.H. Kim, S. Jeong, M. Jeon, H. Lee, A.R. Kim, B. Choi, S.J. Ha, I. Jung, K.J. Yoon, J.E. Park, J.H. Kim, B.J. Kim, E.C. Shin, and S.H. Park. 2023. A virtual memory CD8(+) T cell-originated subset causes alopecia areata through innate-like cytotoxicity. Nat Immunol 24:1308–1317.

Shannon, P., A. Markiel, O. Ozier, N.S. Baliga, J.T. Wang, D. Ramage, N. Amin, B. Schwikowski, and T. Ideker. 2003. Cytoscape: a software environment for integrated models of biomolecular interaction networks. Genome Res 13:2498–2504.

Stuart, T., A. Srivastava, S. Madad, C.A. Lareau, and R. Satija. 2021. Single-cell chromatin state analysis with Signac. Nat Methods 18:1333–1341.

Wang, E.H.C., M. Khosravi-Maharlooei, R.B. Jalili, R. Yu, A. Ghahary, J. Shapiro, and K.J. McElwee. 2015. Transfer of Alopecia Areata to C3H/HeJ Mice Using Cultured Lymph Node-Derived Cells. J Invest Dermatol 135:2530–2532.

Xing, L.Z., Z.P. Dai, A. Jabbari, J.E. Cerise, C.A. Higgins, W.J. Gong, A. de Jong, S. Harel, G.M. DeStefano, L. Rothman, P. Singh, L. Petukhova, J. Mackay-Wiggan, A.M. Christiano, and R. Clynes. 2014. Alopecia areata is driven by cytotoxic T lymphocytes and is reversed by JAK inhibition. Nat Med 20:1043–1049.

Yang, C., D.I. Kwon, M. Kim, S.H. Im, and Y.J. Lee. 2021. Commensal Microbiome Expands Tgammadelta17 Cells in the Lung and Promotes Particulate Matter-Induced Acute Neutrophilia. Front Immunol 12:645741.

Yang, Q., K.R. Safina, K.D.Q. Nguyen, Z.K. Tuong, and N. Borcherding. 2025. scRepertoire 2: Enhanced and efficient toolkit for single-cell immune profiling. PLoS Comput Biol 21:e1012760.

Zhang, Y., T. Liu, C.A. Meyer, J. Eeckhoute, D.S. Johnson, B.E. Bernstein, C. Nusbaum, R.M. Myers, M. Brown, W. Li, and X.S. Liu. 2008. Model-based analysis of ChIP-Seq (MACS). Genome Biol 9:R137.

