## supplementary figures for "Continuous generation of short-lived resident effector CD8 T cells sustains alopecia areata"

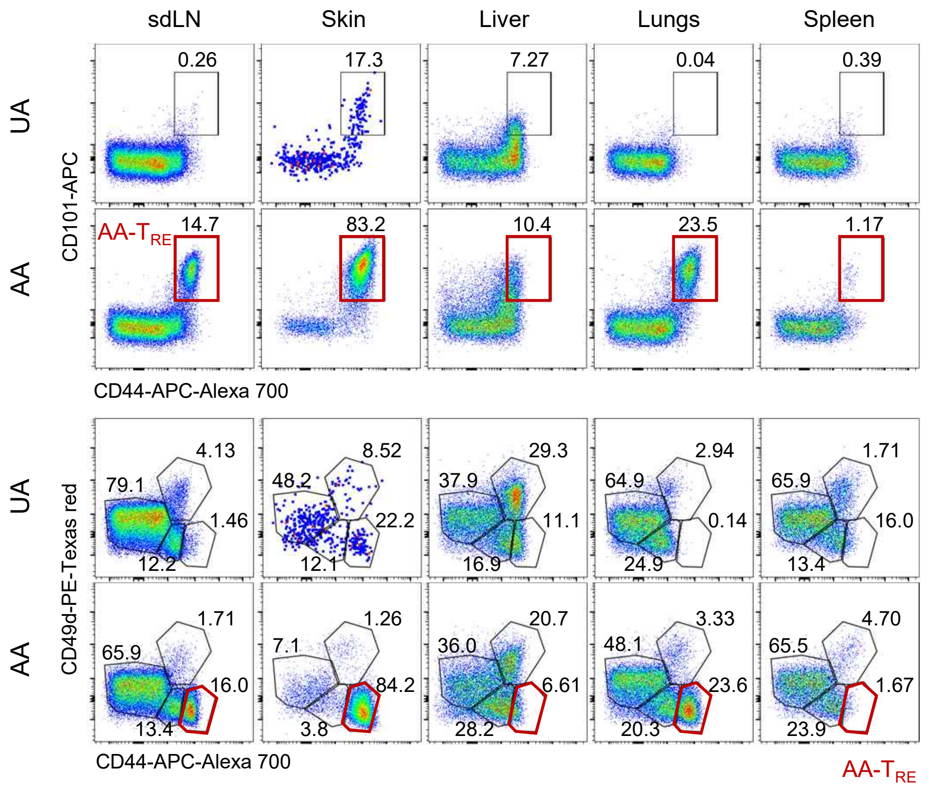


**Supplementary Figure 1. AA is mediated by tissue-resident CD8 T cells.**

Dot plots showing indicated marker expression in UA and AA C3H/HeJ mice in the indicated organs. Representative data from at least 5 independent experiments.

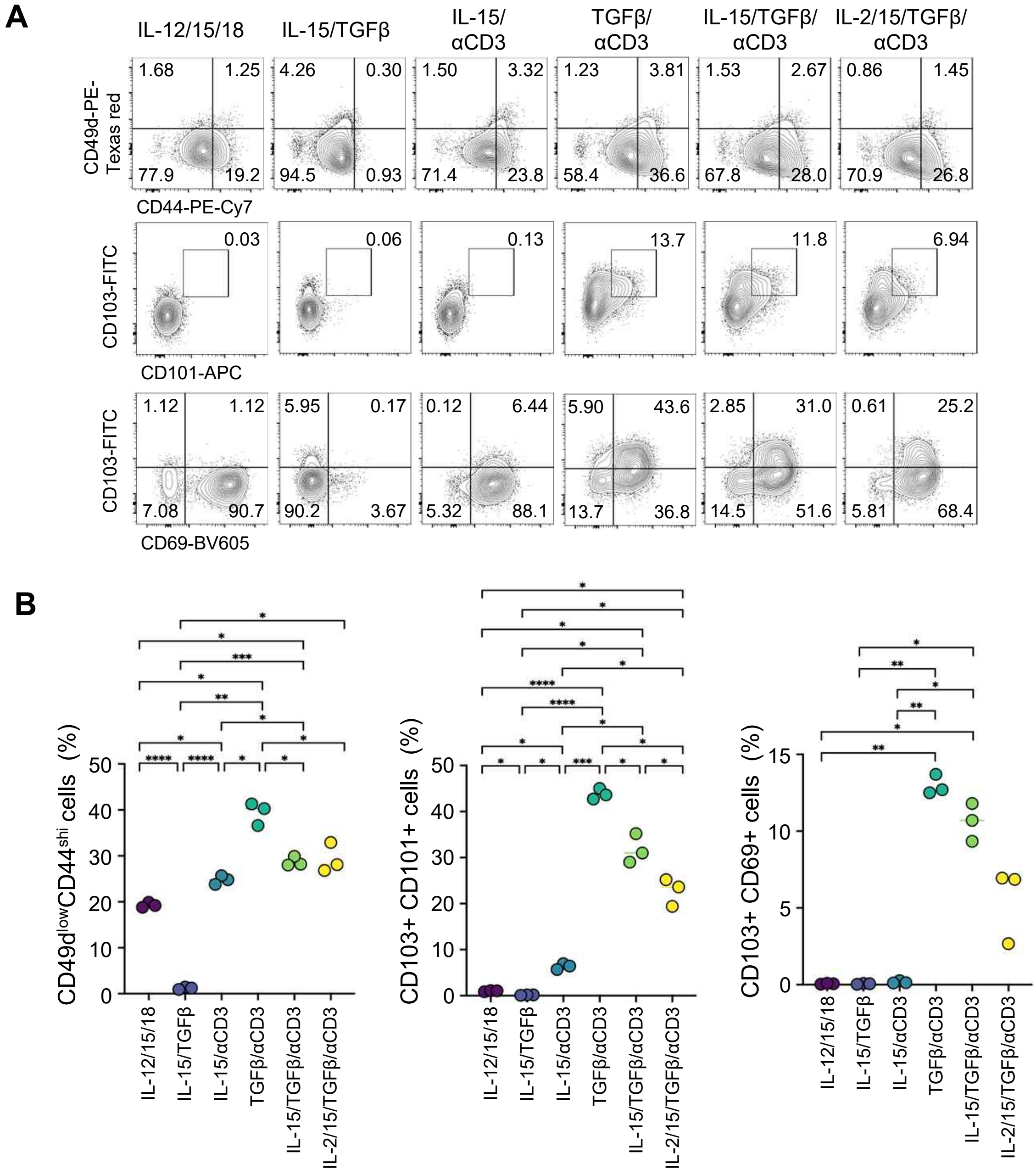


**Supplementary Figure 2. TGF-β and CD3 stimulation drive the differentiation of $T_{RE}$ cells.**

Isolated CD44$^{hi}$ CD8 $T_{MEM}$ cells were stimulated with the indicated cytokines and/or anti-CD3/CD28 Dynabeads for 3 days.

**(A)** Representative flow cytometry plots show expression of indicated markers under each culture condition. Data from three independent experiments are shown.

**(B)** Graphs show statistical analyses.

Numbers indicate frequencies of cells in each quadrant or adjacent regions. Data are mean ± S.D. Statistical analysis was performed using a repeated-measures one-way ANOVA followed by Tukey's multiple comparisons test (B); *p<0.05, **p<0.01, ***p<0.001, ****p<0.0001.

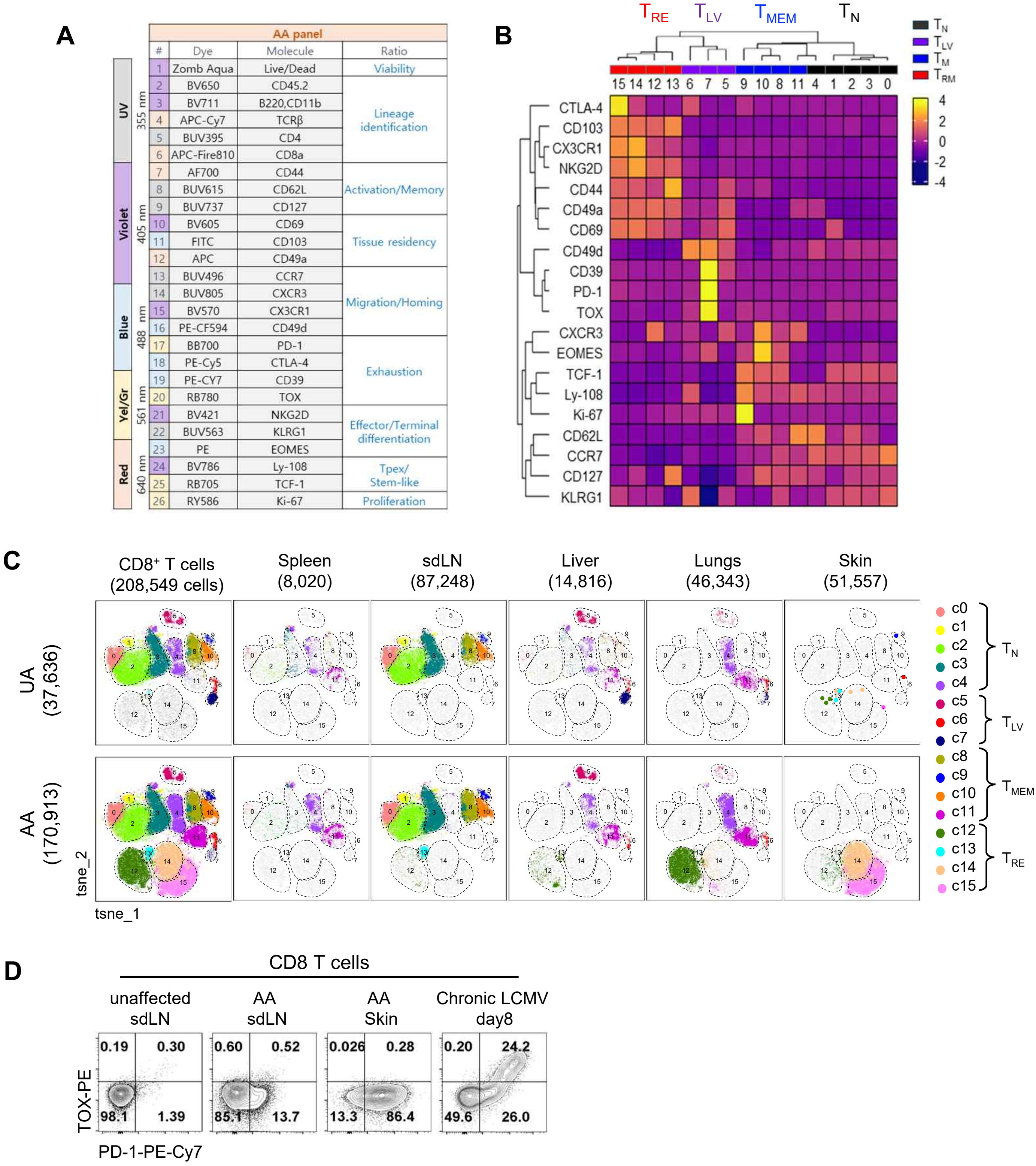


**Supplementary Figure 3. Multiplex analysis of $T_{RE}$ cells.**

**(A-C)** 26-plex cytometry analysis was performed using CD8 T cells isolated from sdLN, skin, liver, lungs and spleen. Antibody panel used for analysis (A). Heat map showing the differential marker expression among $T_N$, $T_{LV}$, $T_{MEM}$, and $T_{RE}$ CD8 T cells in AA (B). t-SNE plots from Figure 2A show the distribution CD8 T cells in indicated organs (C). **(D)** Flow cytometry plots show the expression of TOX and PD-1 in CD8 T cells from C3H/HeJ mice with AA and C57BL/6 mice with chronic LCMV infection.

Representative data from 3 independent experiments are shown. Numbers indicate frequencies of cells in each quadrant or adjacent regions. UA, unaffected; $T_N$, naïve T; $T_{MEM}$, memory T; $T_{LV}$, liver T.

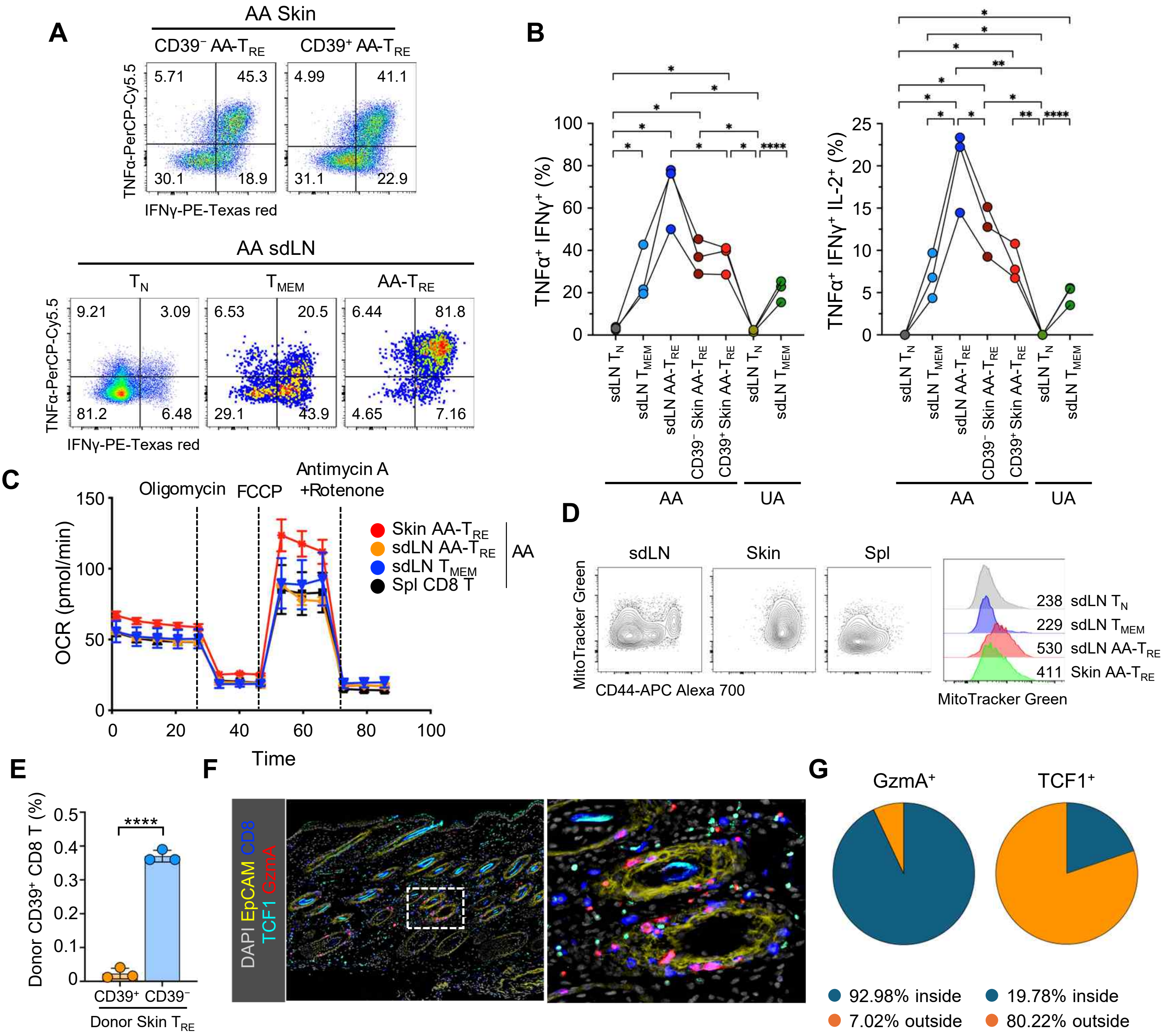


**Supplementary Figure 4. $T_{RE}$ cells are activated T cells.**

**(A-B)** Representative dot plots showing TNF-α and IFN-γ expression in CD8 T cells activated with PMA/ionomycin in the indicated populations (A). Graphs show the frequencies of double (left) and triple (right) cytokine-secreting cells in the indicated cells (n=3) (B). Pooled from 3 independent experiments.

**(C)** OCR analysis of indicated cell populations in AA.

**(D)** Contour plots and histograms comparing MitoTracker Green expression in the indicated organs and cells. Representative data from 3 independent experiments are presented.

**(E)** Graph shows CD39+ donor CD8 T cell frequencies 10 days after transfer (n=3). Results refer to Fig. 2E.

**(F-G)** Immunofluorescence staining using the IBEX technique shows the expression pattern of the indicated molecules in the skin of an AA mouse (F). $GzmA^+$ and $TCF1^+$ cells were quantified using a histocytometric algorithm (G).

Each dot represents an individual mouse. Numbers indicate frequencies of cells in each quadrant or gMFI of cell populations. Data are mean ± S.D. Statistical analysis was performed using a one-way ANOVA with Geisser-Greenhouse correction, followed by a Tukey's multiple-comparisons test (B); and unpaired t-test (E); *p<0.05, **p<0.005, ****p<0.0001. OCR, oxygen consumption rate; FCCP, carbonyl cyanide 4-(trifluoromethoxy) phenylhydrazone; AA, alopecia areata; UA, unaffected; WT, wild-type; sdLN, skin-draining lymph node; $T_N$, naïve T; $T_{MEM}$, memory T; $T_{RE}$, resident effector T.

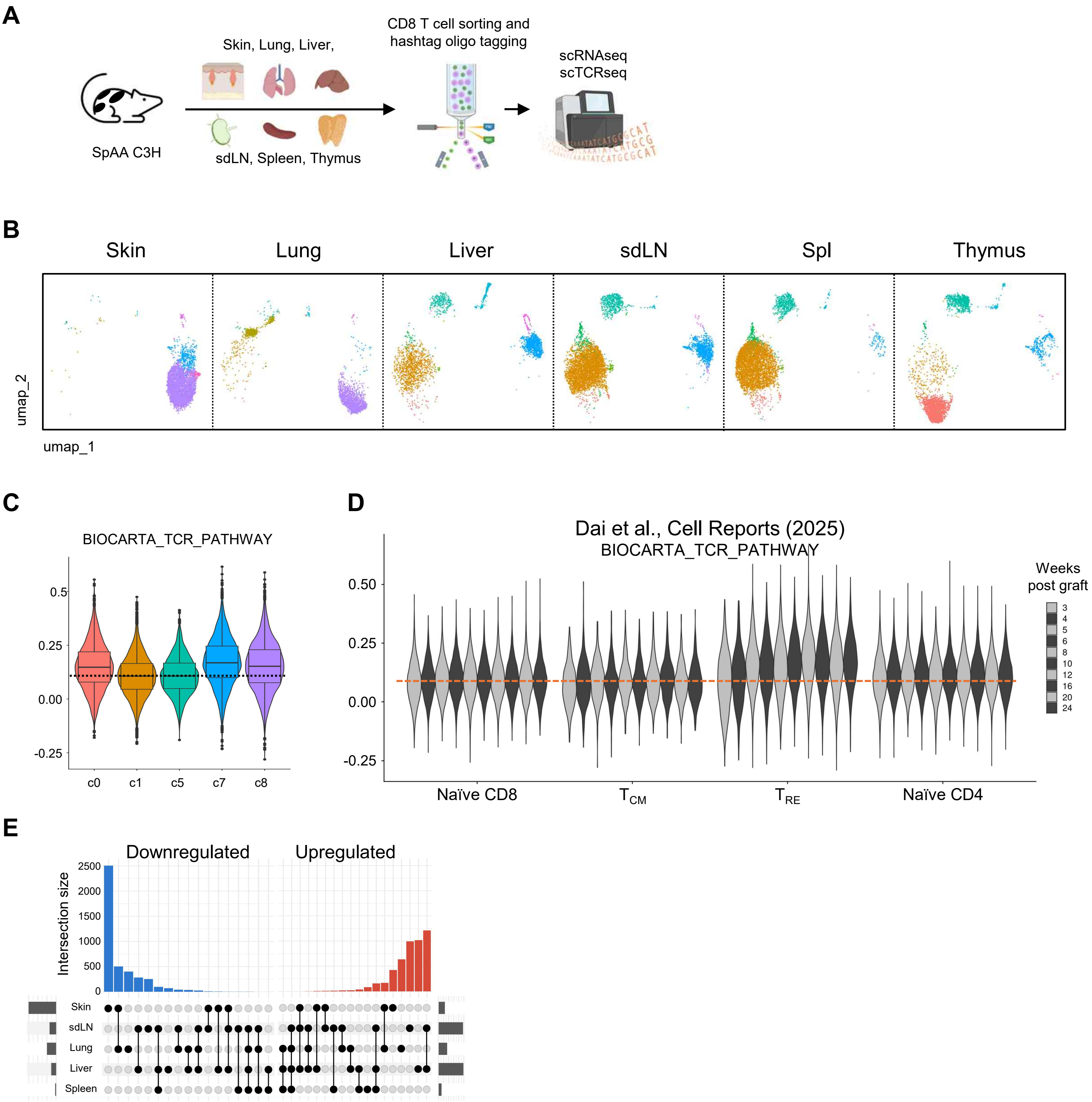


**Supplementary Figure 5. scRNA-seq combined with scTCR-seq analysis of CD8 T cells from AA.**

**(A)** Schematic figure showing the experimental procedure for analyzing CD8 T cells sorted from the indicated organs of C3H/HeJ mice with SpAA. Cells were barcoded with oligonucleotides for demultiplexing before sequencing.

**(B)** UMAP visualization of CD8 T cells separated by organs.

**(C)** Violin plot showing modular scores for the Biocarta TCR pathway in indicated CD8 T cells.

**(D)** Violin plot showing modular scores for the Biocarta TCR pathway in CD8 T cells of C3H/HeJ mice from a public dataset (https://data.mendeley.com/datasets/gxhhxrtr2w/1) that analyzed samples across weeks post-AA induction by skin graft.

**(E)** UpSet plot depicting DEGs of $T_{RE}$ cells across organs.

SpAA, spontaneous alopecia areata.

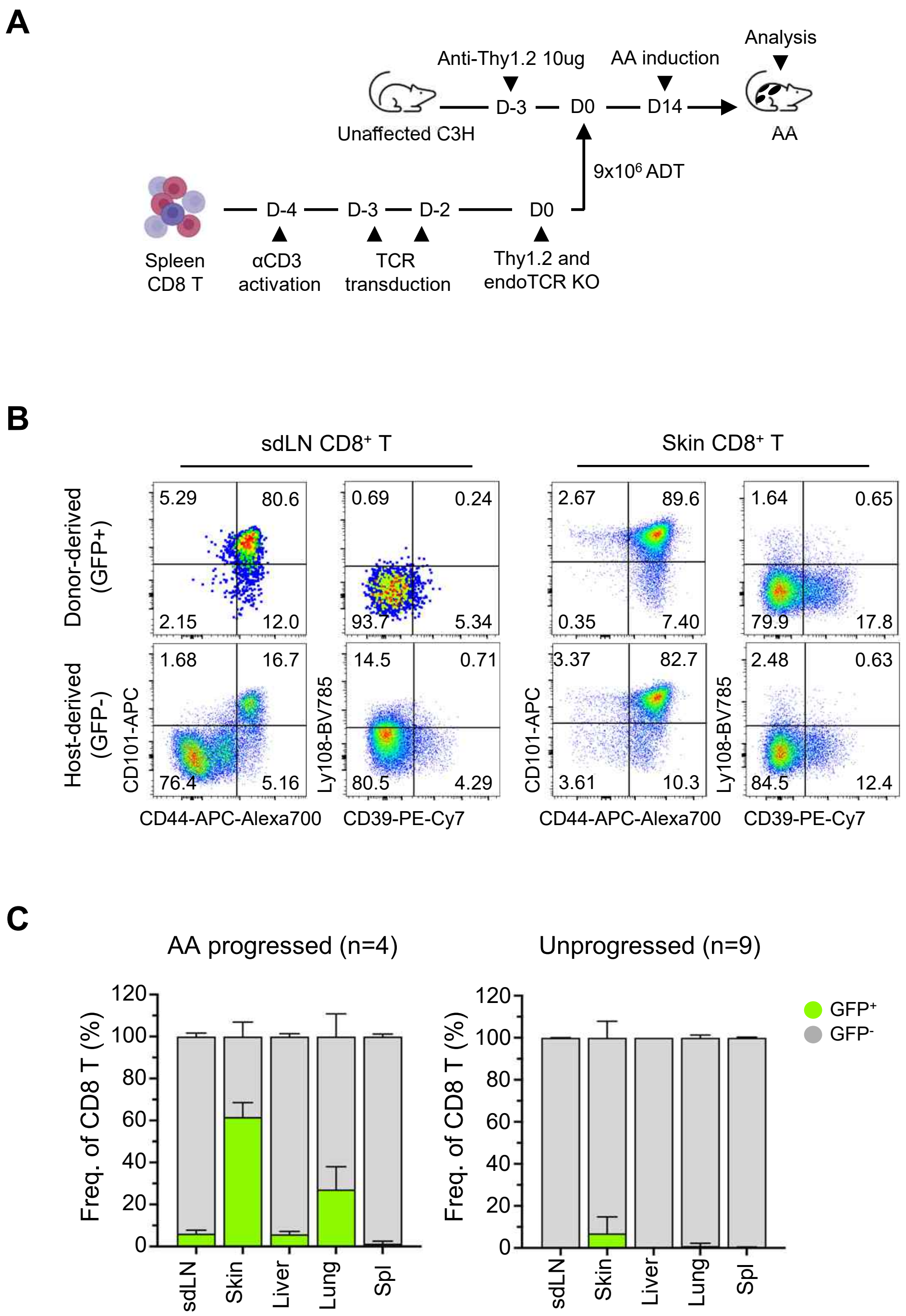


**Supplementary Figure 6. TCRs direct the activation of $T_{RE}$ cells.**

TCR-transduced CD8 T cells were transferred into Thy1-depleted UA hosts that were induced to develop AA and analyzed upon disease induction.

**(A)** Schematic figure showing the experimental scheme.

**(B)** Representative dot plots comparing donor-derived monoclonal (top) and host-derived polyclonal (bottom) CD8 T cells in sdLN and skin of AA-progressed mice.

**(C)** Bar graphs showing proportions of GFP-expressing CD8 T cells in the indicated organs of AA-progressed (n=4) or unprogressed (n=9) mice.

Representative data from 4 independent experiments. Numbers indicate frequencies of cells in each quadrant or adjacent regions. Data are mean ± S.D. UA, unaffected.

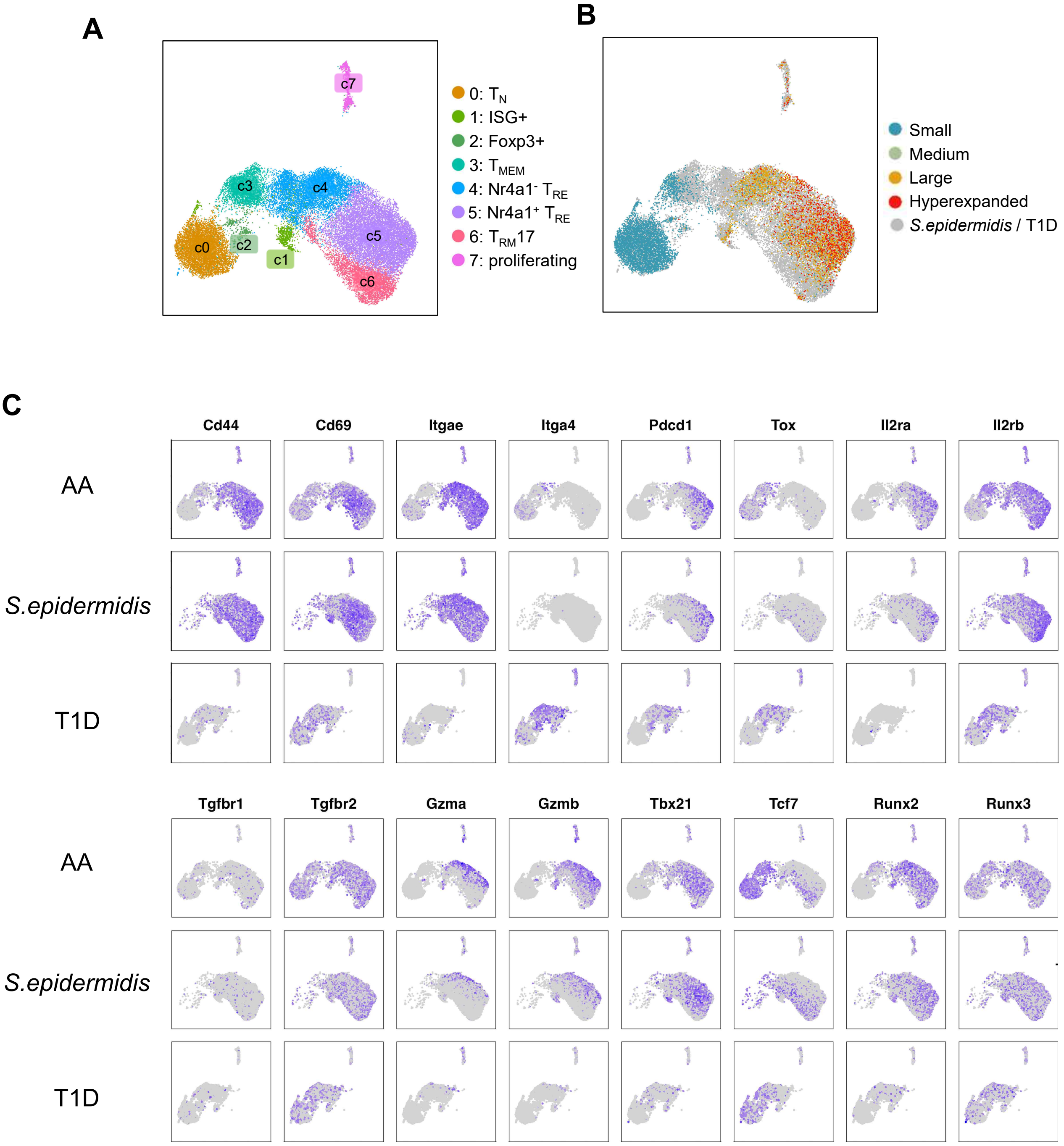


**Supplementary Figure 7. $T_{RE}$ cells exhibit clonal expansion and $T_{RM}$–like transcriptional signatures.**

**(A)** Integrated UMAP of CD8 T cells from AA skin and sdLNs, public *S. epidermidis*–associated skin and lymph node datasets (GSE233389), and autoimmune T1D pLNs (GSE151652).

**(B)** Integrated UMAP displaying clonal expansion of CD8 T cells in AA skin and sdLN; small ≤ 0.001, 0.001< medium ≤ 0.01, 0.01 < large ≤ 0.1, 0.1 < hyperexpanded ≤ 1.

**(C)** Feature plots displaying the expression of indicated genes across different conditions.

ISG, interferon signature gene

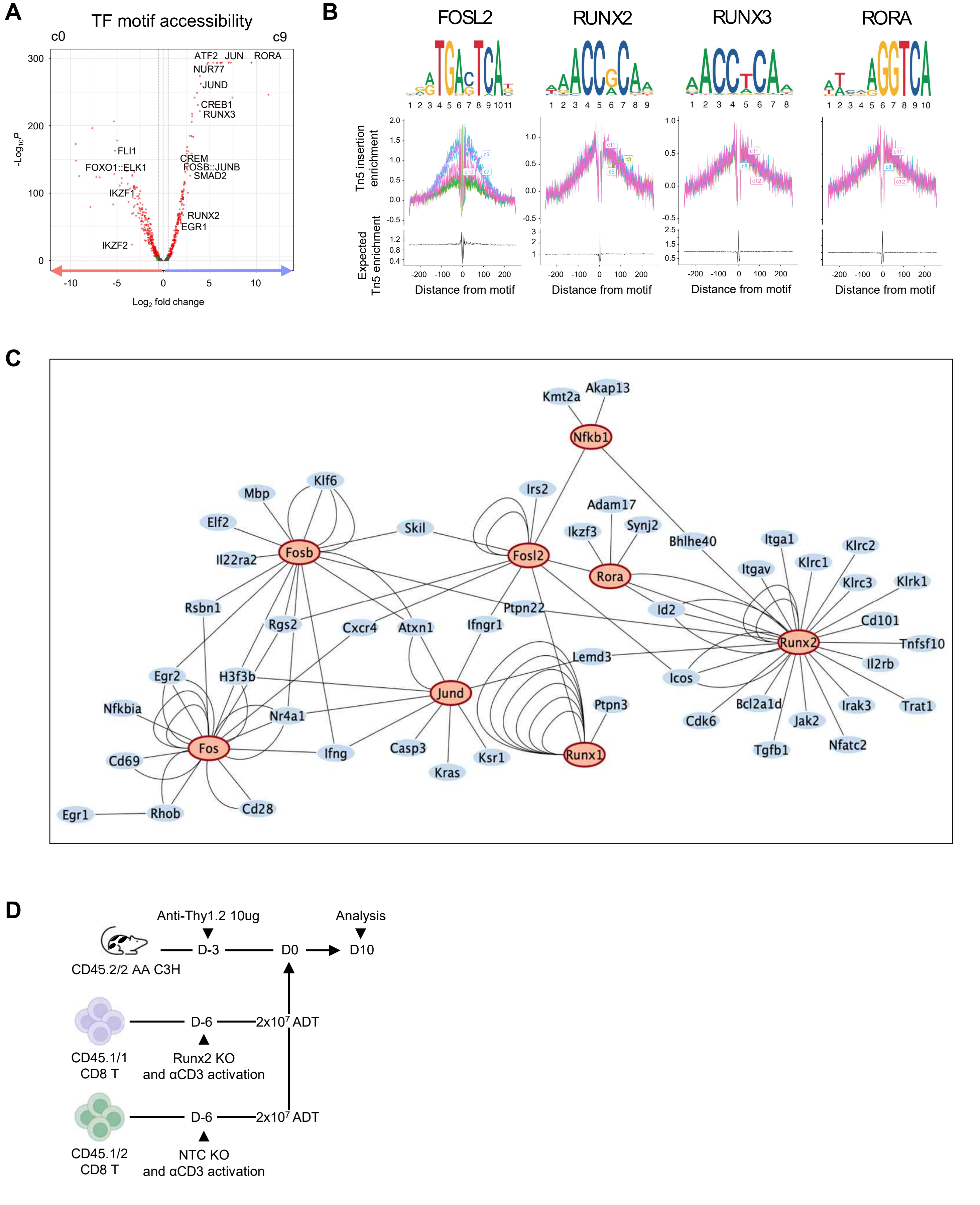


**Supplementary Figure 8. $T_{RE}$ cells exhibit distinct TF activation programs.**

**(A)** Volcano plot shows the differential motif activities between naïve (c0) and $T_{RE}$ (c9) cells in Fig. 4A.

**(B)** Motif position weight sequence and TF footprinting for the indicated TFs are shown.

**(C)** SCENIC+ inferred GRN shows core TF in red, and target genes in blue.

**(D)** Experimental scheme of transferring Runx2 or NTC KO cells into AA hosts

TF, transcription factor; GRN, gene regulatory network.

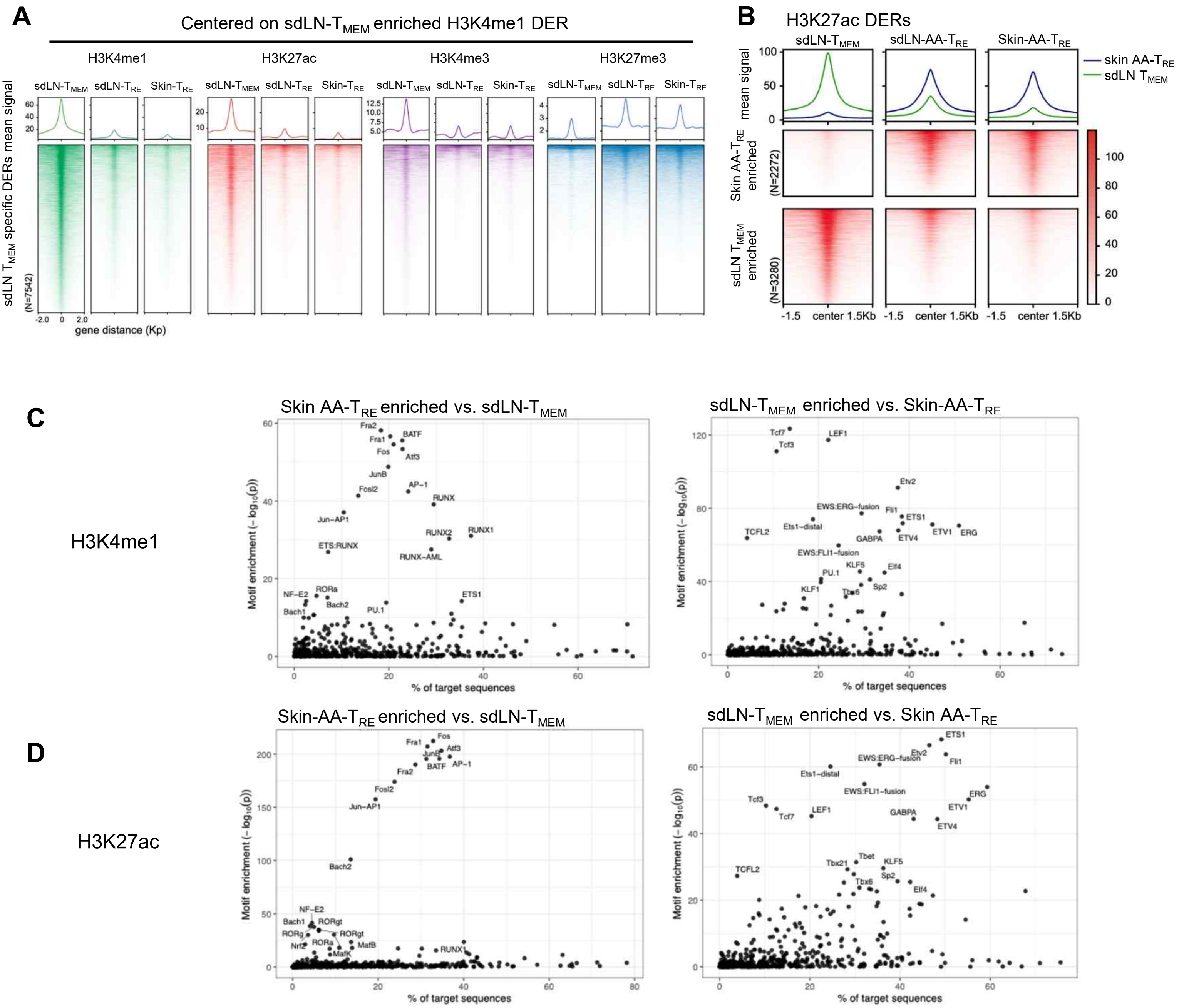


**Supplementary Figure 9. Chromatin enrichment dynamics and TF motif enrichment at H3K4me1 and H3K27ac CUT&Tag-seq DERs.**

**(A)** Aggregated signal profiles (top) and heatmap (bottom) show normalized CUT&Tag-seq signals for multiple histone modifications centered on H3K4me1 DERs enriched in sdLN $T_{MEM}$ cells compared to skin $T_{RE}$ cells.

**(B)** Aggregated signal profiles (top) and heatmap (bottom) show normalized H3K27ac signals centered on H3K27ac DERs comparing sdLN $T_{MEM}$ and skin $T_{RE}$ samples.

**(C)** TF binding motif enrichment analysis performed on upregulated H3K4me1 DERs using HOMER.

**(D)** TF binding motif enrichment analysis performed on upregulated H3K27ac DERs using HOMER.

DER, differentially enriched region.

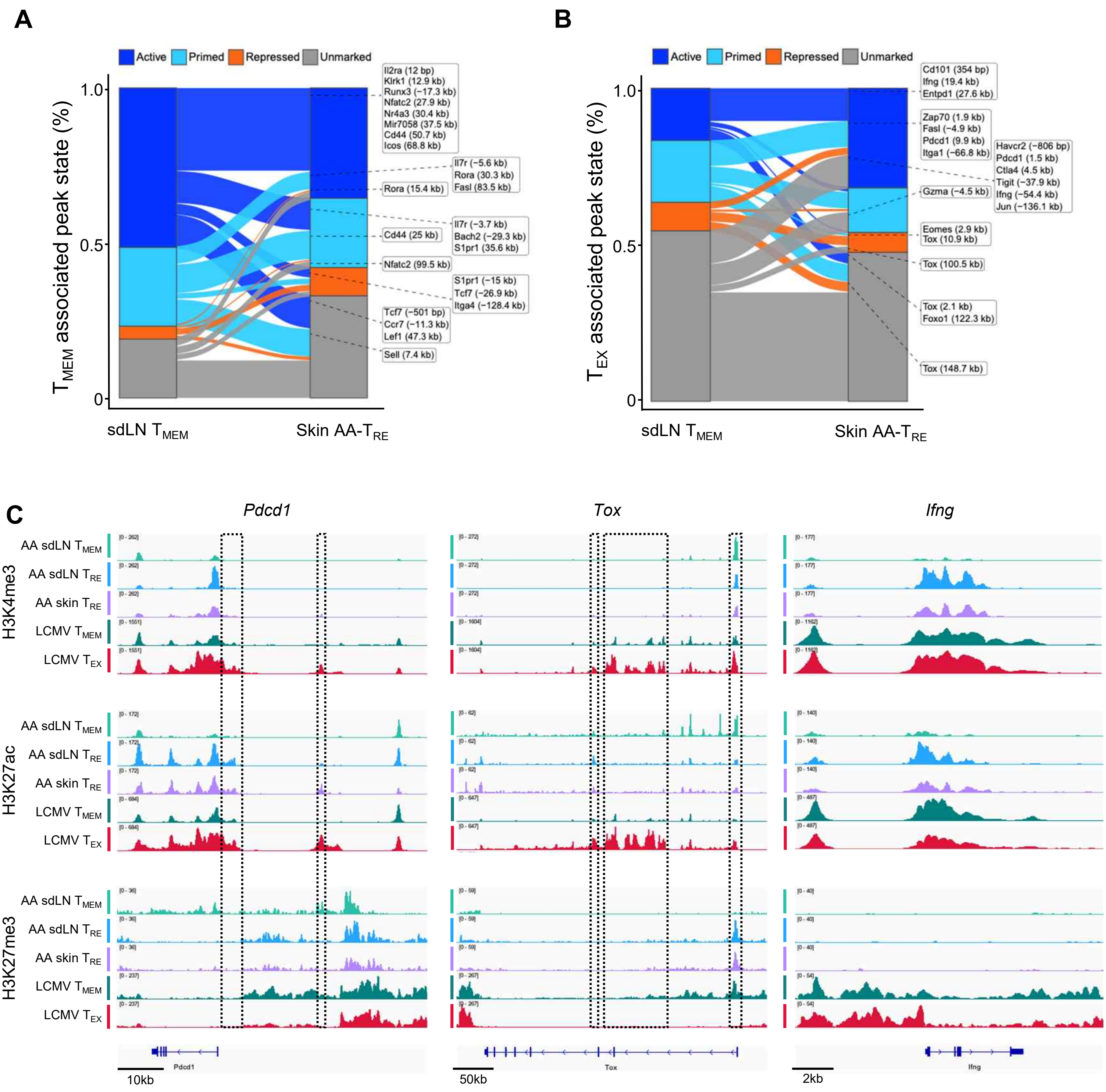


**Supplementary Figure 10. $T_{RE}$ cells partially overlap with $T_{EX}$ signatures in chromatin enrichment dynamics.**
**(A-B)** Alluvial plot displaying the transition of individual $T_{MEM}$ or $T_{EX}$-associated enhancers between chromatin states from sdLN $T_{MEM}$ to skin AA-$T_{RE}$. The width of each flow represents the number of enhancer regions undergoing a given state transition: Active (H3K4me1$^{+}$H3K27ac$^{+}$H3K27me3$^{-}$), Primed (H3k4me1$^{+}$H3K27ac$^{-}$ H3K27me3$^{-}$), Repressed (H3K27ac$^{-}$ H3K27me3$^{+}$), Unmarked (H3K4me1$^{-}$H3K27ac$^{-}$ H3K27me3$^{-}$). Peak locations were defined using published CUT&RUN-seq data (GSE285245).
**(C)** Genome tracks showing H3K4me3, H3K27ac, and H3K27me3 data at the indicated genomic locations. Published CUT&RUN-seq data (GSE285245) were displayed as LCMV $T_{MEM}$ and LCMV $T_{EX}$. Dashed boxes highlight published enhancer regions and additional regions identified in this study.
DER, differentially enriched region.

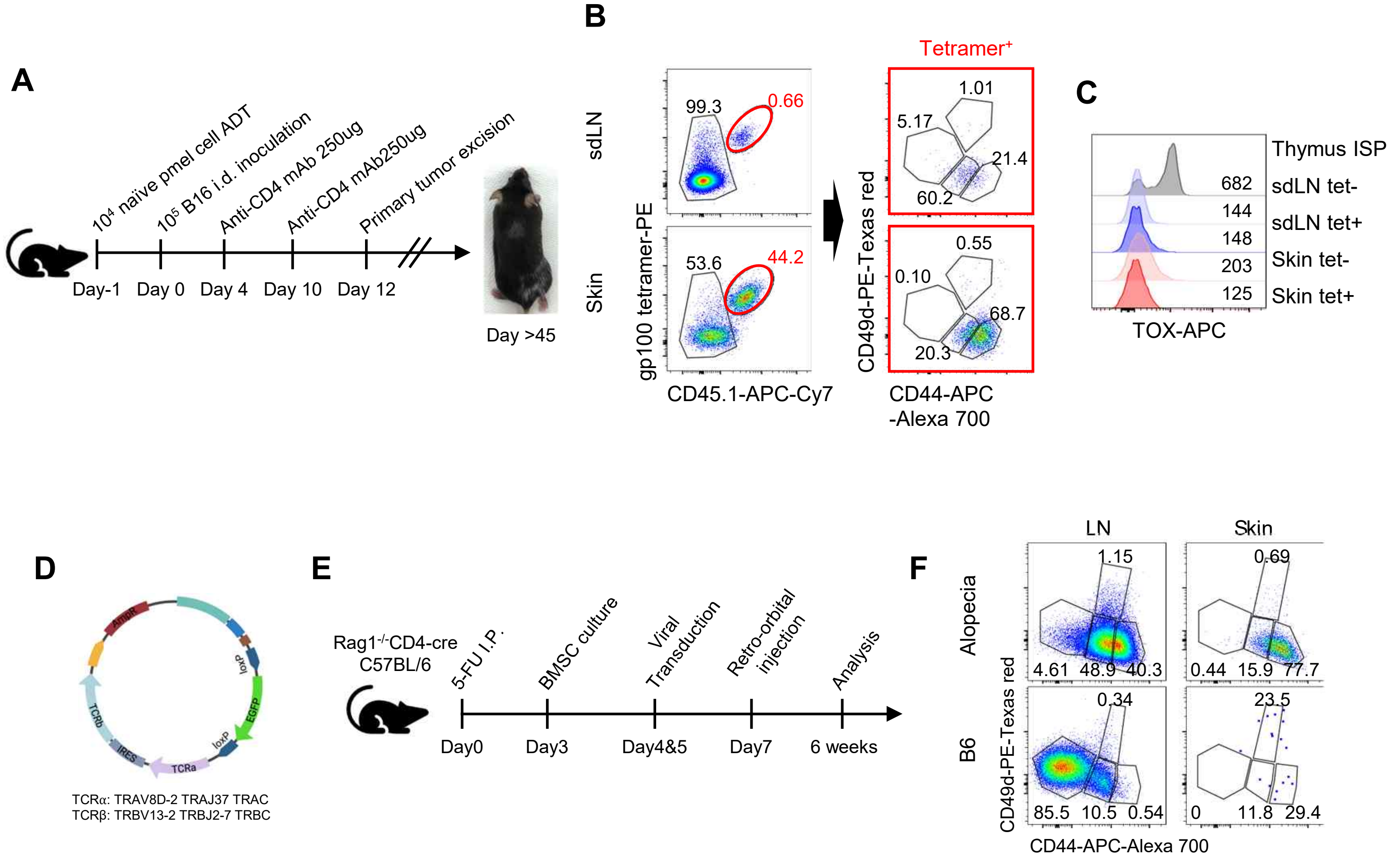


**Supplementary Figure 11. A CD44$^{s\text{-}hi}$ CD49d$^{low}$ phenotype is shared by TCR-dependent skin-resident CD8 T cells.**

**(A-C)** Naïve pmel-1 T cells were adoptively transferred into WT C57BL/6 mice and induced MAV after primary tumor excision. Experimental scheme showing generation of MAV model (A). Representative flow cytometry plots showing indicated marker expression in gp100 tetramer+ CD8 T cells (B). TOX expression in indicated populations of sdLN and skin (C). Representative data from 3 independent experiments.

**(D-F)** B6 mice were induced to develop alopecia using a 1MOG244 TCR retrogenic mouse. Vector map for 1MOG244 TCR (D). Experimental scheme showing the generation of the 1MOG244 retrogenic chimera in C57BL/6 mice (E). Dot plots showing CD49d and CD44 expression in 1MOG244 retrogenic CD8 T cells in the sdLN and skin of alopecia-induced mice (F). Representative from at least 10 independent experiments.

Numbers indicate frequencies or gMFI of cells in adjacent regions. ISP, immature single positive; UA, unaffected; MAV, melanoma-associated vitiligo; BMSC, bone marrow stem cell.
